# Molecular crypsis by pathogenic fungi using human factor H. A numerical model

**DOI:** 10.1101/250662

**Authors:** Stefan Lang, Sebastian Germerodt, Christina Glock, Christine Skerka, Peter F. Zipfel, Stefan Schuster

## Abstract

Molecular mimicry is the formation of specific molecules by microbial pathogens to avoid recognition and attack by the immune system of the host. Several pathogenic Ascomycota and Zygomycota show such a behaviour by utilizing human complement factor H to hide in the blood stream. We call this type of mimicry molecular crypsis. Such a crypsis can reach a point where the immune system can no longer clearly distinguish between self and non-self cells. Thus, a trade-off between attacking disguised pathogens and erroneously attacking host cells has to be made, which can lead to autoreactivity. Based on signalling theory and protein-interaction modelling, we here present a mathematical model of molecular crypsis of pathogenic fungi using the example of *Candida albicans*. We tackle the question whether perfect crypsis is feasible, which would imply that protection of human cells by complement factors would be useless. The model identifies pathogen abundance relative to host cell abundance as the predominant factor influencing successful or unsuccessful molecular crypsis. If pathogen cells gain a (locally) quantitative advantage over host cells, even autoreactivity may occur. Our new model enables insights into the mechanisms of candidiasis-induced sepsis and complement associated autoimmune diseases.

## Introduction

Crypsis and mimicry are wide-spread phenomena in biology. They are used to deceive predators, prey, hosts or other interaction partners. Most commonly they are used as defensive strategies like imitation of harmful species by harmless species (Batesian mimicry) or imitation of a dominant element of the environment (cryptic mimesis). Aggressive mimicry refers to the observation that some predators use camouflage not to be recognized by their prey; mimicry can also be useful for reproduction, for example, in egg mimicry realized by some birds (Zabka and Tembrock, 1986; Pasteur, 1982; Vane-Wright, 1976). The effect of mimicry can be understood by analogies from human society. For example, uniforms may be misused by civilians, as was impressively described in the theatre play “The Captain of Koepenick” by Carl Zuckmayer (Zuckmayer, 1932) and in the movies “The Sting” and “Catch Me If You Can”.

Crypsis and mimicry require three entities: a model (in the case of crypsis this is the environment) which is imitated by a mimic to deceive a dupe (also called operator). Usually mimicry systems are discussed with respect to animals and plants (Dittrich et al., 1993; Huheey, 1988; Holen and Johnstone, 2004), but they also occur in the realm of pathogenic micro-organisms. The mimicked trait is often a chemical compound (for example, when predatory animals roll on the grass to hide their odour from potential prey) rather than a visual feature. Of course, in the case of micro-organisms, visual features are irrelevant anyway. Damian (1964) introduced the term “molecular mimicry” to describe antigen sharing between host and parasites. Later Damian (1989) loosened its definition to the microbial production of “similar or shared molecular structures”, preventing host response.

Also nowadays, molecular mimicry research focuses mainly on the adaptive immune system (Tsonis and Dwivedi, 2008; Blank et al., 2007; Cusick et al., 2012; Oldstone, 1998). In this paper an example of molecular crypsis by pathogenic fungi so as to deceive innate immunity is studied, notably the recruitment of a special human immune regulatory protein, the complement factor H (FH). This is performed by various fungal pathogens such as *Candida albicans* (Meri et al., 2002), *Staphylococcus aureus* (Sharp et al., 2012) or *Borrelia burgdorferi* (Haupt et al., 2007; Kraiczy et al., 2001; Hellwage et al., 2001; Blom et al., 2009). We perform our analysis using the example of *C. albicans* although the utilization of human complement regulatory proteins to avoid complement attack is a general phenomenon and the analysis is easily adaptable to be applied to other pathogens.

FH together with complement component 3 (C3) are key factors of the complement system (Zipfel and Skerka, 2015). Both are continuously present at high concentrations in the blood. C3 is cleaved to C3b, which can bind covalently to the surface of host and pathogen cells (Sim et al., 1981; Pangburn and Müller-Eberhard, 1984; Lambris et al., 2008). Together with complement factor B and activated by factor D, C3b can form the C3b convertase (C3bBb), again converting C3 into C3b, so that a local amplification of C3b generation by a positive feedback loop occurs (see Fig. 1).

**Figure 1:**
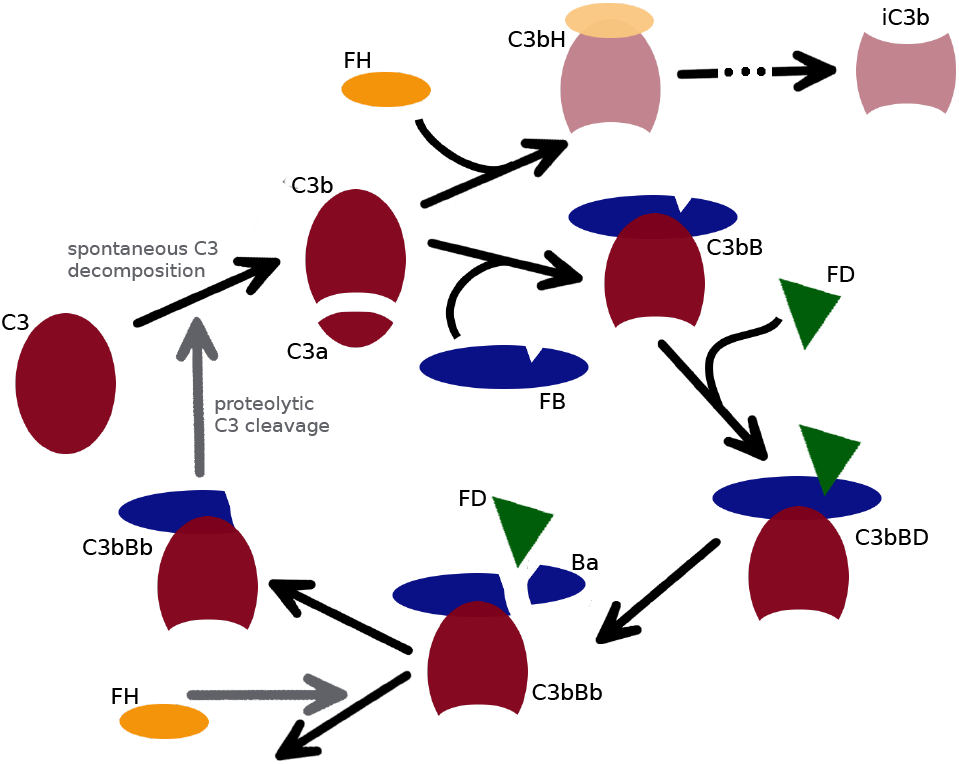
Generalized scheme of C3b amplification and regulation. C3b activation can occur in fluid phase or on surface. Under physiological conditions, C3b amplification in fluid phase is prevented by FH. On cell surfaces, amplification may occur, depending on whether or not FH can be acquired there. If FH cannot be acquired in sufficient amounts, there is a steady but slow conversion of C3 to C3b initiating amplification. C3b then associates with factor B (FB) to form the C3 proconvertase (C3bB), which is activated by factor D, resulting in the active C3 convertase (C3bBb). C3bBb in turn is able to proteolytically convert C3 into C3b at a high rate, starting the loop again and thus acting as an amplifier of C3b production. The amplification loop can be controlled by FH at two stages. Firstly, FH competes with FB for C3b binding, acting as a cofactor for C3b degradation to iC3b. Secondly FH is able to accelerate the decay of C3bBb, reducing proteolytic C3b generation.

The accumulation of C3b on the cell surface is called opsonization. The deposited C3b on the surface acts as a signal for phagocytes, like macrophages, to remove the tagged cells. Additionally opsonization with C3b has various downstream effects on the innate, as well as the adaptive immune system. For example, it mediates the formation of a terminal complement complex (TCC) at a high opsonization state, effectively perforating the cell membrane and triggering lysis of the cell (Lambris et al., 2008).

C3b binds to all types of surfaces, i.e. self and nonself (Sim et al., 1981). The discrimination between self and non-self is accomplished by FH and other complement regulators (Zipfel and Skerka, 2009). FH can bind to glycosaminogens on the host surface, like heparan sulfates and to surface deposited C3b. Once bound to the cell surface, FH prevents C3b amplification and is able to inactivate surface-bound C3b. For damaged host cells, C-reactive protein (CRP) can prevent binding of FH to the cell surface, thus maintaining opsonization (Zipfel and Skerka, 2009; Lambris et al., 2008). The main expression site of FH and C3 is the liver, but local expression can be accomplished by other cell lines as well, for example, (in the case of FH) endothelial, epithelial or muscle cells (Ferreira et al., 2010; Uhlén et al., 2015).

Molecular crypsis enters the scene in that several pathogenic fungi and micro-organisms bind FH or other complement regulators by, for example, the complement regulators acquiring surface proteins (CRASP), like pH regulated antigen (Pra1) of *C. albicans* and other proteins (Meri et al., 2002; Sharp et al., 2012; Haupt et al., 2007; Kraiczy et al., 2001; Hellwage et al., 2001; Blom et al., 2009). Depending on the effectiveness by which pathogens recruit FH and the physiological parameters of the host, this could prevent or reduce C3b opsonization of pathogens to some degree. From a macrophage’s “viewpoint”, this may increase the similarity between host and pathogen cells. In consequence, the host may have to decide on a trade-off between erroneously attacking own cells, possibly inducing autoreactivity, and the effectiveness by which it is able to clear mimetic pathogens from the blood (see Fig. 2). This trade-off may not always be easy to find, which might be related to the presence of two different FH alleles in the human population, one leading to a higher risk of autoreactivity than the other (Hageman et al., 2005; Hummert et al., 2017). This illustrates the role of complement as a ’double-edged swor? (Zipfel et al., 2013; Jacob et al., 2010; Liu et al., 2011) where the immune system may erroneously attack own cells and, on the other hand, pathogens may mimic host cells as an evasion strategy (see Dühring et al. (2015) for alternative evasion strategies).

**Figure 2:**
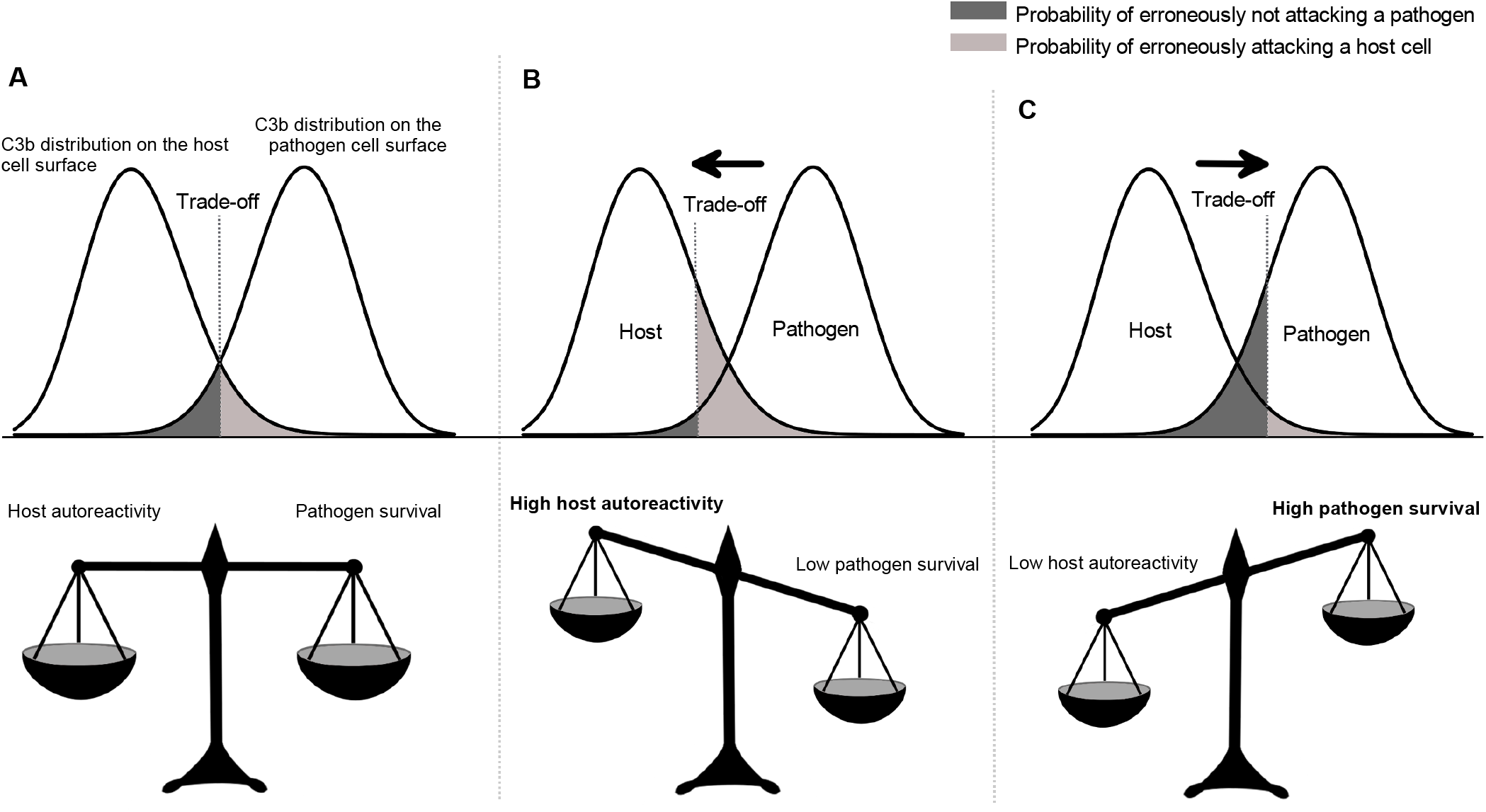
Example C3b distributions with differently weighted trade-off. This figure illustrates the attack decision problem the host faces for different C3b distributions on host and pathogen cell surfaces. All cells with a higher amount of C3b bound to the cell surface than the threshold will be attacked. (A) Equally weighted trade-off; the probability of erroneously attacking a host cell is equal to the probability of erroneously not attacking a pathogen cell. (B) Trade-off weighted in favour of the effectiveness in clearing mimetic pathogens. (C) Trade-off weighted in favour of low autorcactivity.

One may have different opinions on which phenomenon occurs. Pasteur (1982) doubts relationships of molecular mimicry to organismic mimicry because in the former case there is “no sensory perception involved”. Nevertheless in the case of molecular mimicry antigens are bound by antibodies which activate Fc-receptors on the surface of phagocytes. Also in the case of (unsuccessful) molecular crypsis, various receptors are activated. Activation of those receptors triggers specific signalling pathways, leading to a specific response of the phagocytes. Thus we adopt the view that sensory perception, although on a molecular level, is involved and molecular mimicry should be analysed like any other mimicry system. Furthermore we consider FH recruitment to prevent C3b opsonization as a new type of molecular mimicry with the difference that instead of mimicking a specific antigen (actively transmit a self signal), pathogens prevent deposition of specific opsonins to camouflage in the environment of the blood-stream (actively inhibit a non-self signal).

Accordingly, we classify this system as molecular cryptic mimesis (molecular crypsis), involving the unusual case that the environment is not neutral to the dupe. This is because the environment is a multicellular organism to which the dupe (phagocytes) belongs. When pathogens use camouflage, they make, for phagocytes, the decision of whether or not to attack cells very hard because it may happen that own cells are attacked.

Host-pathogen interactions have been successfully analysed by mathematical modelling (Anderson and May, 1982; Hummert et al., 2013; Dühring et al., 2017; Becker et al., 2018) In particular, mathematical models describing mimicry by using, for example, methods from game theory, have been presented earlier for higher organisms, either in a general context (Holen and Johnstone, 2004; Speed and Ruxton, 2010; Huheey, 1988) or for specific types of mimicry (Huheey, 1976; Speed, 1993). In contrast, models describing molecular mimicry of bacteria or fungi are rare. Our paper is aimed at filling this gap and extending theoretical mimicry research to pathogenic micro-organisms including fungi. This helps us provide an alternative view 011 a wide variety of complement associated autoimmune diseases, including the ambiguous or paradoxical role complement plays in the pathogenesis of sepsis (Markiewski et al., 2008).

The question arises whether the pathogen is capable of perfect mimicry (camouflage). In that situation, the human immune system would not be able to distinguish between human (own) cells and pathogen cells. Then, however, protection of human cells by complement factors would be useless. On the other hand, complement factors are successfully used. Our study is aimed at investigating how the immune system can resolve that paradox.

## Materials and methods

We start by analysing the attack-decision problem the host faces. Based on a single trait, which is here opsonization with C3b, phagocytes have to decide whether or not to attack a single cell, being either a host cell or a pathogen. As a result of this decision two errors may arise: erroneously attacking a host cell and erroneously not attacking a pathogen. Using an approach to study mimicry in higher organisms (Holen and Johnstone, 2004; Speed and Ruxton, 2010), we can quantify each of the errors, depending on the encounter rate and the pathogenity of the intruder and the aggressiveness of the host’s immune system (above which threshold of C3b opsonization the host attacks). Based on those errors, their respective probabilities and their severity, we use a population-based evolutionary model to outline the key parameters which influence molecular crypsis in a general context.

In the second part, we examine opsonization states of an individual infection scenario. This is done by adapting the quantitative model of human complement system described in Zewde et al. (2016) to account for FH recruitment of pathogenic microbes. For the parameter values in the model, experiments were performed and complemented by data proposed in Zewde et al. (2016) (Tables A1 and A2). Our adapted model is used to predict physiological C3b opsonization states for the two microbial species *Candida albicans* and *Escherichia coli*, each in comparison to human erythrocytes. It is worth mentioning that *E. coli* is able to perform molecular crypsis, utilizing for example the C4b-binđing protein, which may also have C3b-đegrađing function (Wooster et al., 2006). However, to our knowledge, it is not able to utilize FH. Based on surface area deviations within cell populations we further generate C3b distributions for a single infection with one of the two microbes, which are then used to analyze possible strategies the host could adopt to react to molecular crypsis of pathogens.

### Computing distributions of surface-bound FH

The binding reaction of FH to a binding site on the cell surface can be described in an approximative way by the following reaction equation:

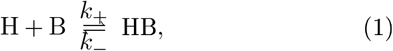

where *k*_+_ and *k*_–_ are rate constants. Let *H* and *HB* denote the number of free and bound FH molecules, respectively, and *B* the number of free binding sites. Then, for the mean dissociation constant *K_d_* of all molecules binding FH on the cell surface, we have

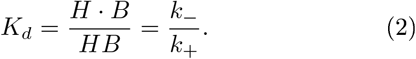

Defining *n* as the total number of all binding sites on pathogen surfaces in the medium, i.e.

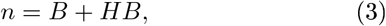

it follows for the dissociation constant

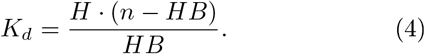

This can be rewritten as

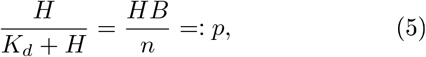

where we define the variable *p* for the fraction of occupied binding sites, or, the probability that a single binding site is occupied. For the immune system, it is of interest how many of all binding sites are occupied. This number is here denoted by *k*. The binomial distribution applies because that corresponds to the number of *k*-combinations from a given set of *n* elements:

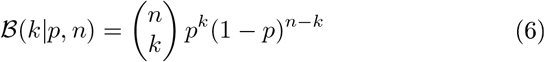

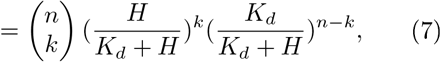

Note that *n* – *k* binding sites are not occupied. The expected value *μ* and the standard deviation *σ* of this distribution are

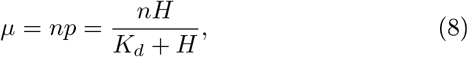

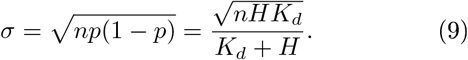

Note that we can generally assume large *n* on the host side, but also on the pathogen side if cell numbers are high enough. For large *n* the binomial distribution may converge to a Poisson distribution (*p* → 0) or a normal distribution, as depicted in Fig. 2.

### Defining the attack decision

Approximation of the attack decision follows the theoretical work of Holen and Johnstone (2004) on mimicry systems and the work of Wiley (1994) on signal perception in biological systems.

Given a particular C3b opsonization state *k*, the decision of attacking can be made by the host using any function, for example a sigmoid, where all trait values *k* above a certain threshold *t* will be attacked. For simplicity we will assume a step function. This implies that once a cell is opsonized sufficiently, there will be no chance evading phagocytosis or lysis.

Given a threshold *t* the two errors arising can be quantified using the cumulative distribution functions 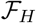 and 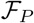 of the C3b distribution of host and pathogens. This distribution depends conversely on the distribution of surface bound FH derived above and is quantified with real-world data in the protein-interaction model later. That is, 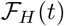 is the probability of erroneously attacking a host cell and 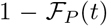 is the probability of erroneously not attacking a pathogen.

If there is no further information on the two signal distributions, signalling theory suggests the optimal way of discriminating the signals to be minimisation of a linear combination of these two errors, where the weights depend on the encounter probabilities, which are proportional to the respective cell numbers (Wiley, 1994; Holen and Johnstone, 2004). It should be considered that in the case of infection by a pathogen, the weighting factors may be time-dependent because the cell numbers of pathogens vary. Thus, it would be beneficial to adjust the threshold during inflammation. It has indeed been observed that autoreactivity and inflammation are correlated (Semin Hematol. 2017 Jan;54(1):33-38. doi: 10.1053/j.seminhematol.2016.10.003. Lymphocyte generation and population homeostasis throughout life. Yanes RE, Gustafson CE, Weyanđ CM, Goronzy JJ.; Ref.). We consider this effect in that the optimal threshold depends on pathogen abundance.

Moreover, in systems where the attack decision implies some benefit or cost, depending on whether it was right or erroneous, the probabilities of the two errors should additionally be weighted by the benefit in correctly attacking a pathogen and the costs arising by erroneously attacking an own cell (Wiley, 1994). The more virulent a pathogen is, the higher is the benefit of attack.

Taking all together, in the presented model, it is beneficial for phagocytes to choose a threshold *t*\*, where the difference of the probability of attacking a pathogen, weighted by the benefit *B* and the probability of attacking an own cell weighted by the cost *C* is maximized:

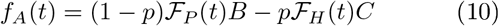

Note that in this formula, we assume an infection with a single pathogen, so the encounter probability of the pathogen can be approximated by (1 – *p*) with *p* being the encounter probability of the host. A more exact model should not only include the frequency-but also the density dependency.

Assuming a normally distributed trait value with standard deviation *σ* (Note that for high receptor numbers the binomial C3b distribution converges to a normal distribution) Holen and Johnstone (2004) identified the optimal attack threshold for this type of fitness function to be:

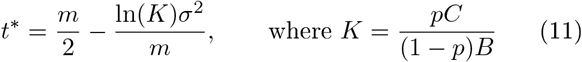

In this formula m is the distance in the trait value space of both species. So for *m* = 0 the species are only separable by their standard deviation, which is assumed to equal σ for both species in this case. In our model *m* is the difference in C3b opsonization of host and pathogen, where the host is assumed to have *m* = 0 in a physiological state. Strictly speaking, *m* is not a constant value in our model. For example C3b amplification on pathogen surfaces may influence the opsonization state of the host. This effect is examined by the protein interaction model in detail.

We see that the optimal threshold depends substantially on the dissimilarity *m*, where higher similarity results in a higher (more aggressive) threshold, but this is dampened by the mimetic load *K*, which increases with low probability of encountering a pathogen or low benefit to-cost-ratio of attacking that pathogen (low pathogenity).

### The population based evolutionary model

For the evolutionary model we assume that the C3b distribution of a given species is defined by the ability of FH surface acquisition alone. This implies that in this case all species below the FH threshold concentration 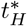 are attacked (because we assume those have a C3b opsonization above the “real” threshold regarding C3b). We next define the maximum possible binding sites the cells can produce (restricted due to limited energy and/or space), called *n_max_*. Furthermore we define the relative energetic cost *c* ∈ [0,1] to produce these binding sites, compared to the whole available energy. The overall payoff *u* of a species *S* is the product of the energy available for growth and maintenance after receptor production and its survival probability. This energy is, for simplicity’s sake, assumed to depend linearly on *n*:

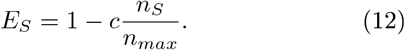

The payoff for the pathogen then reads

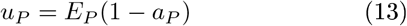

where we use the abbreviation 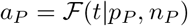, which is the pathogen’s probability of being attacked, given threshold *t*.

The payoff of the host *H* will depend on the attack decision:

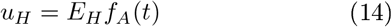

If we assume that both the pathogen and the host can alter their amount of receptors and their dissociation constant and the host decides which threshold it should assign to a given pathogen (i.e. based on its degree of mimetic resemblance and on the pathogenity in general) strategies of pathogen cells are a pair (*n, K_d_*), with *n* ∈ *N* = {0, 1,…, *n_max_*}, while host strategies are a triple (*n, t, K_d_*) with *n, t* ∈ *N*. Thus, the host has one degree of freedom more than the pathogen. Note that by decreasing the threshold for pathogenic cells which can bind FH very effectively the host can avoid autoimmunity, but has to rely on other mechanisms of immune defense than the complement system.

Optimization of the parameters was done using an evolutionary algorithm (Beyer and Schwefel, 2002)starting from random populations of hosts and pathogens .

### The protein-interaction model

To investigate C3b opsonization in a single infection scenario, we have adapted the models presented by Zewde et al. (2016) and Korotaevskiy et al. (2009) to account for FH surface acquisition of pathogens. Our adapted model focuses on the initial part of complement activation. Thus we did not consider stabilization of the C3bBb complex with properdin or formation of the TCC. Our model of initiation of the alternative pathway of complement has 28 species and 38 reactions. The parameter values of FH binding to host and pathogen surfaces were obtained by own experiments. Red blood cell data was taken from the combined NHANES datasets from 2001 to 2014 (Centers for Disease Control and Prevention (CDC). National Center for Health Statistics (NCHS), 2014). The remaining parameter values were taken from Zewde et al. (2016). Altogether the parameter values were chosen so as to provide realistic estimates of the conditions encountered in the blood stream.

We implemented the model using the COmplex PAthway Simulator, COPASI (Hoops et al., 2006). Units of the model are micromole for amount of substance, litre for volume and milliseconds for time. C3b amplification may occur in blood plasma, serum or on surfaces. In serum it is activated by a spontaneous tick-over reaction of C3 into C3_*H*_2_*O*_. As C3_*H*_2_*O*_ has similar properties to C3b, we did not distinguish between C3_*H*_2_*O*_ and C3b in the model, but between fluid and surface bound molecular species (denoted by the prefixes f and b respectively). C3b will then associate with factor B (FB) to form the C3 proconvertase (C3bB), which is activated by factor D, resulting in the active C3 convertase (C3bBb). C3bBb in turn is able to convert more C3 into C3b, starting the loop again (see Fig. 1).

In contrast to the spontaneous cleavage, the proteolytic cleavage of C3 into C3b results in a reactive intermediate called nascent C3b (nC3b), which can bind covalently to cell surfaces before it associates with water to form C3b (Sim et al., 1981). Fig. 3 depicts the processes around the binding of C3b on cell surfaces. The processes shown in Fig. 1 are part of these and are depicted there in more detail.

**Figure 3:**
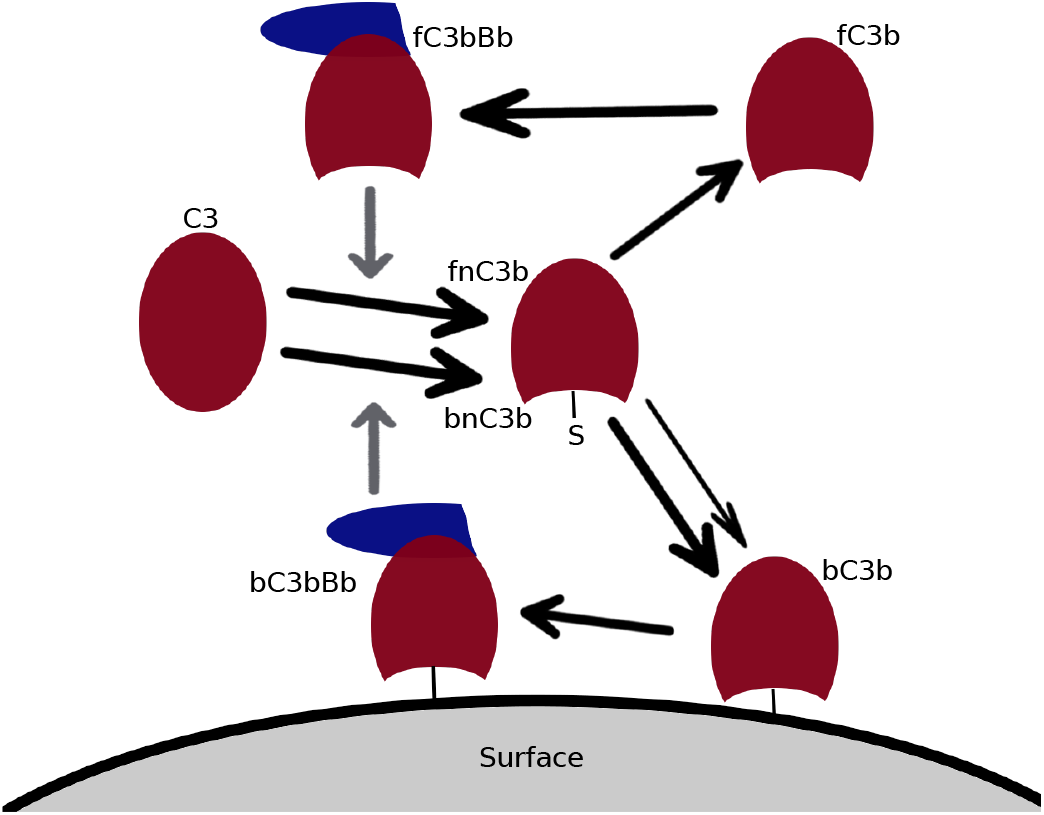
Scheme of C3b surface binding. Proteolytic cleavage of C3 by either fluid (denoted by the prefix f) or surface bound (denoted by the prefix b) C3 convertase (fC3bBb or bC3bBb) results in a reactive intermediate (fnC3b or bnC3b) which is able to covalently attach to surfaces due to an exposed thioester bond. While chemically there is no difference between fnC3b and bnC3b, we distinguish them in the model, because we assume a higher binding rate of bnC3b to the originating surface (see text and A.1.5).

If nC3b encounters a surface, the same positive feedback as in fluid may occur, where the tick-over (basal rate of spontaneous decay) is replaced by the rate of attachment of nC3b to host or pathogen surfaces 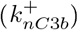. Proteolytic cleavage of C3 by the C3 convertase assembled on surfaces (bC3bBb) will again result in nascent C3b (bnC3b). Although fnC3b and bnC3b are chemically identical, the dynamics of bnC3b surface binding in an ODE system will differ from those of fnC3b, as was already pointed out by Zewde et al. (2016). This is because the local concentration of binding sites surrounding a bC3bBb unit will differ from those of fnC3b, as was already pointed out by Zewde et al. (2016). This is because the local concentration of binding sites surrounding a bC3bBb unit will differ from the global concentration (B_C3b_) due to spatial effects which should not be neglected (see section A. 1.5 and Fig. A7). This will effectively increase the rate of bnC3b binding to their originating surfaces, compared to other surfaces or fnC3b binding.

A diagram of all reactions, complement concentrations and kinetic rate constants used, as well as the complete ODE model, can be found in the Appendix (see Fig. A2, tables A1 and A2 and section A.4).

Generally, we distinguish two cases, one where the pathogen species is not able to acquire FH on the cell surface (represented by *E. coli*) and one where the species is able to acquire FH (represented by *C. albicans* which can bind FH by the surface molecule Pra1). The host (represented by erythrocytes) can always acquire FH by heparan sulfates (HS) on the cell surface. In the case that FH can be acquired, we assume the same amount of FH binding sites compared to C3b binding sites, which is the whole surface (maximum number). This assumption is supported by the results of population based simulations.

As was already mentioned in the section on Dynamics of C3b opsonization, we included some spatial aspect into the ODE model, by assuming a higher affinity of surface derived nascent C3b to the originating surface. Derivation of the scaling factor and simulations not considering this scaling can be found in the Appendix (section A. 1.5 and Fig. A7).

Complement factors B, D and I were fixed at their physiological concentrations (Table A1). C3a and iC3b were remained fixed at zero concentration. C3 and FH were modelled with an explicit inflow and outflow limited only by blood flow. Inflows and outflows were chosen such that the steady state corresponds to the physiological concentrations of C3 and FH. This is to prevent unrealistic behaviour, where especially C3 consumption grows infinitely due to the positive feedback loop in C3b production at high surface concentrations. A detailed description of assumptions regarding C3 and FH dynamics and sensitivity of the model with respect to these assumptions can be found in the Appendix (Figs. A3, A5 and A6).

During simulations, host cells were inoculated into the medium first and the system was given enough time to approach a steady state. In the case that no FH and C3 inflow occurs, the system cannot approach the physiological steady state because the initial concentrations of C3 and FH are not sufficient to cover all cells simultaneously. In this case we gave the system several C3 and FH pulses corresponding to physiological concentrations until steady state was reached (see Fig. A4 for example runs).

After 1000 seconds, a pathogen is assumed to enter the bloodstream, where the concentrations of pathogenic C3b and FH (if suitable) binding sites were set to the concentrations calculated from the pathogen count defined. Final concentrations of all relevant chemical species were obtained after additional 800 seconds have passed and the system again had enough time to reach steady state (half an hour of simulation in total). Because of the pathogen arrival, we had to use fixed time points for measurements and could not use steady-state information directly.

If distributions were available (in particular, of the erythrocyte count as well as erythrocyte and pathogen surface area, see Fig. A1), 1000 values were sampled from those distributions to obtain C3b distributions for host and pathogen cells at specified pathogen concentrations. The optimal attack threshold in terms of signalling theory was computed for distributions as described in the previous section.

## Results

### Population based evolutionary model

Fig 4 shows the results of the population based model if there is no restriction on parameter values. Remember that the evolutionary model uses the simplification that if FH is bound, no C3b amplification can occur. Therefore all cells with a lower amount of FH bound to the surface than the threshold are attacked in this model (gray shaded area), instead of all cells above a certain C3b threshold.

**Figure 4:**
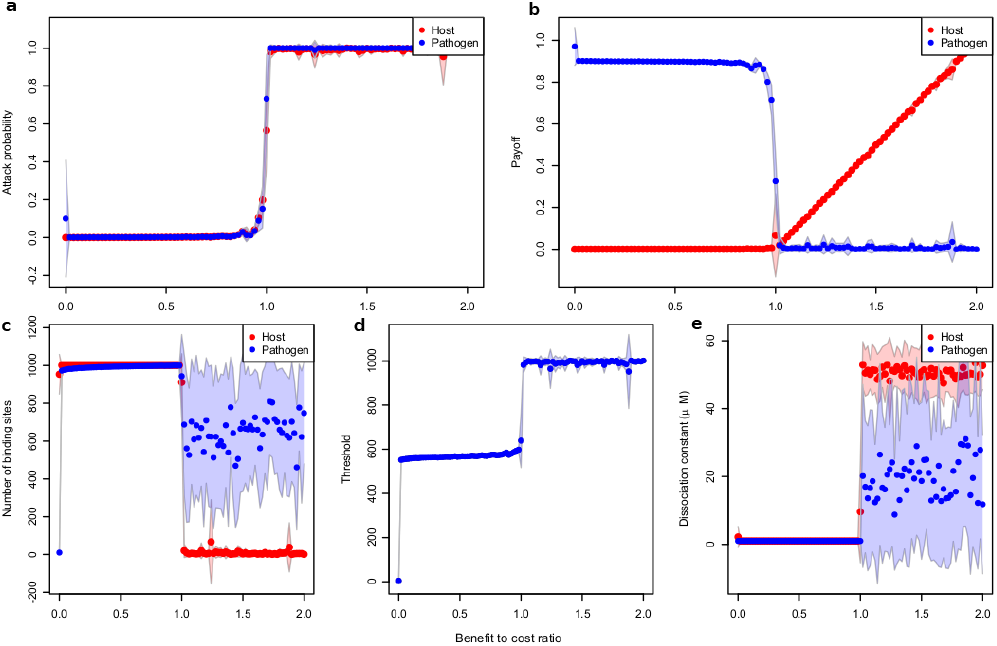
Plot of essential quantities as functions of the benefit-to-cost ratio, computed by numerical optimization. a) Mean probabilities of being attacked by phagocytes for pathogen and host cells, b) Payoff, c) Optimal number of binding sites, d) Classification threshold, e) Dissociation constant. Standard deviation is indicated. Maximum number of binding sites n_max_, 1001; metabolic cost c, 0.1 for both the host and pathogens, 20 runs. FH concentration was set to 1.61 μM.

If there is no benefit in attacking the pathogen at all (in which case it is no pathogen), no cells are attacked (threshold 0) and no binding sites are produced, yielding the highest payoff (1 for pathogen and host). As soon as there is some benefit in attacking the pathogen, both the pathogen and host will start producing their maximum number of binding sites and minimizing the dissociation constant for FH. In consequence, the FH and therefore the C3b distributions on the cell surface will be almost identical and the host cannot distinguish between self and non-self. This leads to a scenario where a lower number of cells are attacked if the benefit in attacking a pathogen is smaller than the cost of attacking an own cell, but all cells are attacked if it is the other way around. Note that when all cells are attacked, there is a random drift in the dissociation constant and the payoff of the host will increase linearly with *r*. Hosts save the cost of producing binding sites altogether in this case, while pathogens do not.

Fig. 5 shows the fitness landscapes of pathogens with fixed host parameters and unlimited (A) or restricted (B) binding site production of the pathogen. In both of the scenarios, there is one local fitness maximum if no binding sites are produced at all and one global maximum at perfect (possible) resemblance. Nevertheless we can observe that the global maximum at the best possible resemblance tends towards the local one as pathogens are less able to obtain perfect resemblance (similar results are observed when restricting for example metabolic energy gain or binding effectiveness).

**Figure 5:**
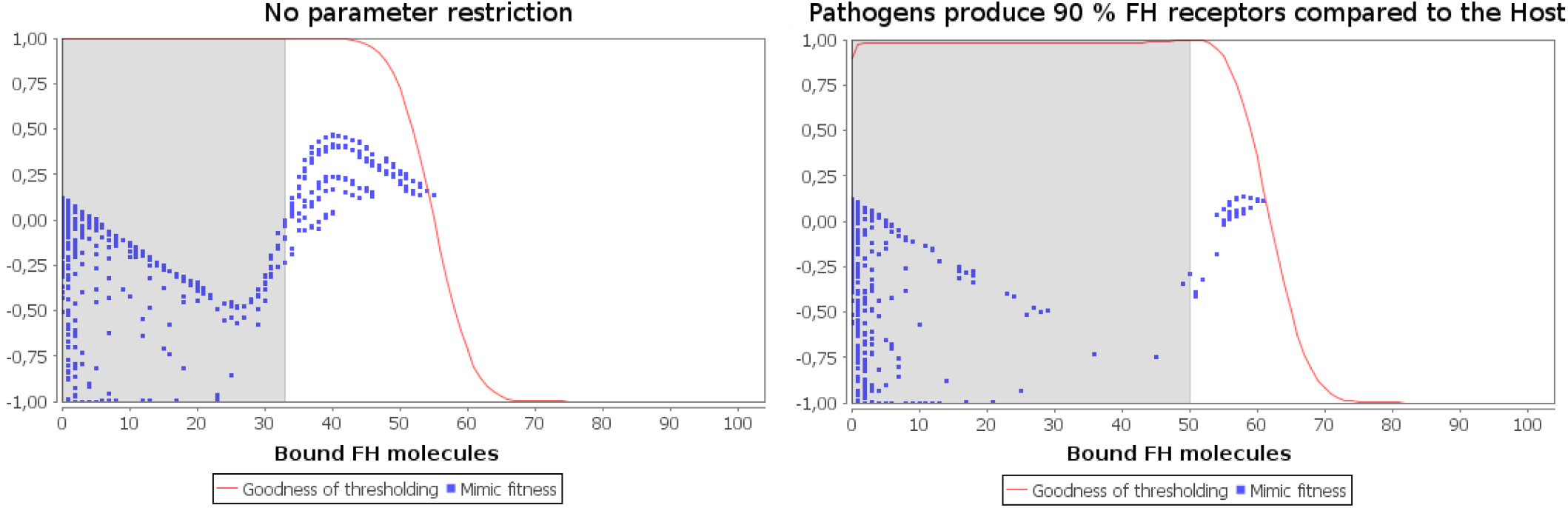
Plot of the fitness landscape of pathogens with fixed host parameters. Equal pathogen and host concentrations were used. Each blue dot represents one individual with a certain investment into the number of binding sites produced (x-axis location) and a certain fitness (y-axis location, determined by the attack probability solely). Host investment into number of binding sites is arbitrarily fixed to 40 % (left) and 60 % (right). In the right subfigure, pathogens are restricted to produce a maximum of 90 % of host binding sites.

### *C. albicans* opsonization

In the following, C3b opsonization means all C3b derived chemical species bound on the surface of host or pathogen. This is the sum of bC3b, bC3bB and bC3bBb concentrations.

Fig. 6 shows the mean C3b opsonization per cell and sample distributions at given pathogen concentrations if factor H can be acquired by the fungus. For low *C. albicans* concentrations compared to erythrocyte concentrations (less than 10^8^ *C. albicans* cells per litre, 1: 50000), host cells have less than one molecule C3b per cell bound on average, while *C. albicans* opsonization is about 10^6^ molecules per cell, corresponding to the whole surface opsonized. Host opsonization then increases until a *C. albicans* concentration of 5 · 10^11^ cells per litre is reached (ratio of 1:10 *C. albicans* vs. erythrocyte cells).

**Figure 6:**
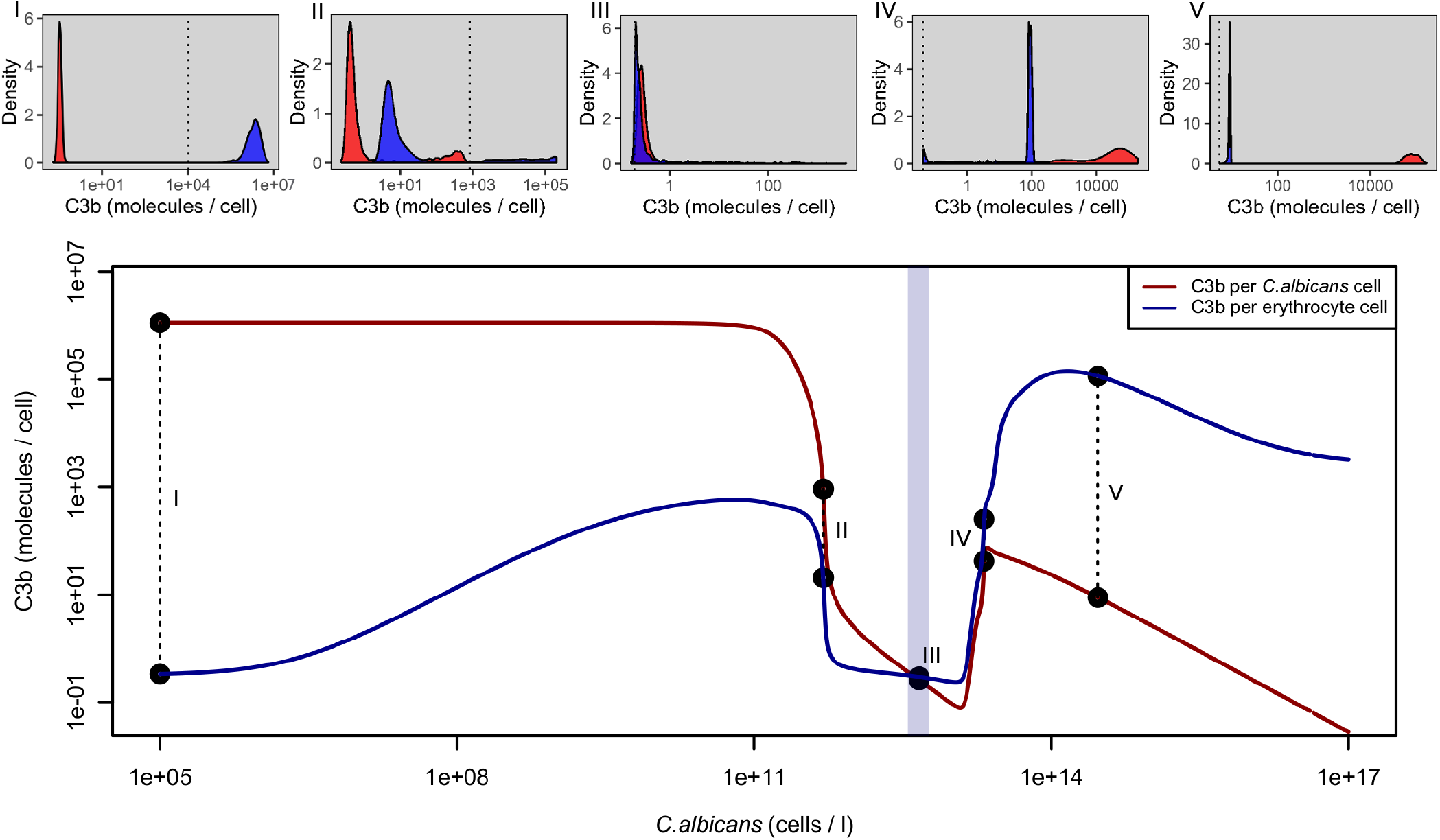
Computed C3b opsonization per cell if factor H can be acquired on the cell surface of the pathogen. Upper panels, distributions of opsonization for different *C. albicans* concentrations. Microbial cell densities for subfigures (I)-(V) can be seen from the corresponding points in the lower panel (see also text). The dotted vertical line in each subfigure represents the mean value of the optimal threshold interval to distinguish the two signals. Lower panel, mean opsonization on host and pathogen surfaces as functions of pathogen concentration, double logarithmic plot. Erythrocyte counts used are indicated by the light blue shaded area (mean 4.64 · 10^12^ erythrocytes per litre, low: 1st percentile, high: 99th percentile, see Fig. A1). C3b opsonization is similar to the case where no FH can be aquired up to a *C. albicans* density of approx. 5 · 10^11^ (i.e., erythrocytes are still 10 times more abundant). Beyond this point crypsis is successful with low opsonization in general. If the pathogen concentration increases even more, competition for factor H dominates and opsonization of both species increases with higher opsonization of host cells until the point where the inflow of C3 is not sufficient to maintain opsonization and C3b decreases again, notably faster for pathogen cells. Separability of the two signals for macrophages is only possible for low pathogen densities. At high pathogen load, the host C3b distribution is similar to the pathogen’s C3b distribution at low pathogen load (subfigures (I) and (V)).

After this point, opsonization of both pathogen and host cells sharply decreases, reaching the opsonization state of the host as if no pathogen were present. The distribution at this point shows that the predicted attack threshold is high, implying very low autoreactivity but effective crypsis of a great deal of pathogens. This behaviour continues until *C. albicans* reaches nearly the same concentration as the erythrocytes (4.64 · 10^12^). Actually the turning point (the intersection point of the red and blue curves in Fig. 6) is slightly less than the mean erythrocyte concentration. Beyond this point, the signals become clearly inseparable and the predicted threshold is very low, meaning the attack of the majority of host and pathogen cells.

For even higher *C. albicans* concentrations, we observe competition for FH (see Fig. A3) and opsonization increases for both host and pathogen cells, while increasing faster for host cells. At a *C. albicans* concentration of about 2.1 · 10^13^ (ratio of 5:1 *C. albicans* vs. erythrocyte cells) opsonization starts decreasing again for the pathogen, while opsonization of host cells still increases. After a *C. albicans* concentration of about 2.96 · 10^14^ is reached (ratio of 65:1 *C. albicans* vs. erythrocyte cells), the C3b distributions of host and pathogen are basically transposed compared to the case where pathogens are absent or present in low numbers in the medium (subfigures (I) and (V) of Fig. 6).

In the case of molecular crypsis of pathogens, the model suggests three discriminable regimes of opsonization, mainly depending on the concentration of pathogens in the blood (see Fig. 7). In the first regime (low pathogen concentration), molecular crypsis is not effective and the host is able to clearly discriminate between self and nonself. In the second regime (medium pathogen concentration), molecular crypsis is successful and opsonization of host and pathogen is low. In the third regime (high pathogen concentration) complement is hyper-active on self, leading to autoreactivity. High autoreactivity due to high pathogen cell density is doubtless not beneficial for the host but may also not be beneficial for the pathogen (Damian, 1964). It is an interesting question in this context if there are mechanisms of pathogens to avoid too high concentrations in the host system.

**Figure 7:**
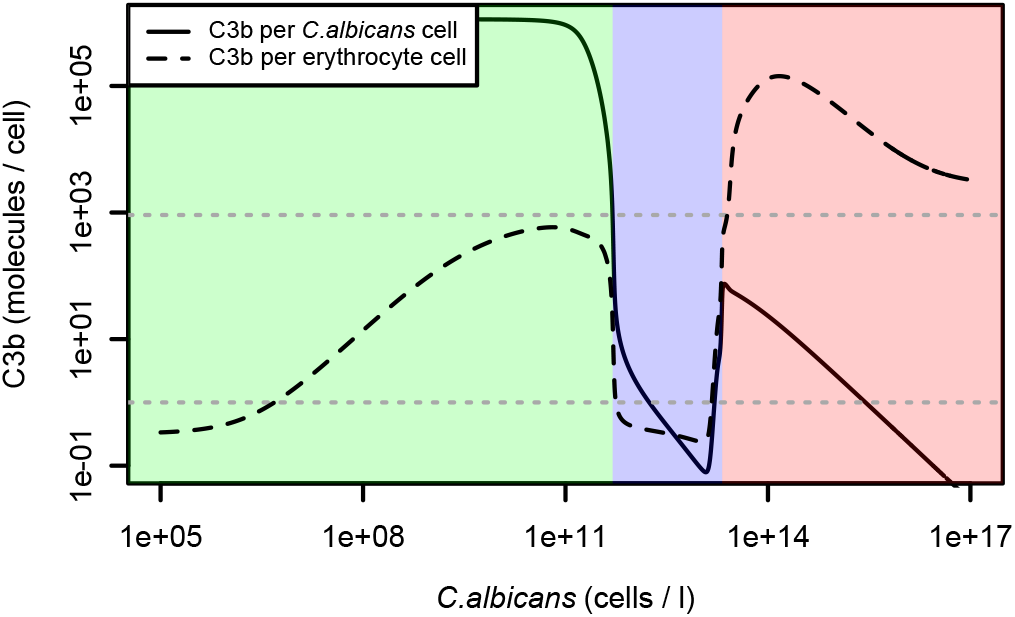
Mean opsonization of erythrocytes and *C. albicans* cells with different regimes and possible attack thresholds indicated. Same data as in Fig. 6. The two dotted horizontal lines represent possible attack thresholds. Upper line, optimal threshold derived from simulations on *E. coli*: lower line, already a single C3b molecule is sensed on the surface. Cells with values above those lines would be attacked. Green, non-successful crypsis: blue, regime of successful crypsis: red. autoreactivity. These regimes are indicated for the upper attack threshold. Using the lower threshold, autoreactivity may occur even for low pathogen concentrations. For any of these thresholds and for any other threshold tested (see subfigures of Fig. 6). host opsonization drastically increases for high pathogen concentrations, making autoreactivity nearly unavoidable.

### *E. coli* opsonization

Fig. 8 shows the mean C3b opsonization per cell and sample distributions at given pathogen concentrations if no factor H can be acquired by the microbes. For low *E. coli* concentrations compared to erythrocyte concentrations (less than 10^8^ *E. coli* cells per litre, ratio of 1: 50000), host cells have less than one molecule of C3b per cell bound on average, while *E. coli* opsonization is about 10^5^ molecules per cell, corresponding to the whole surface opsonized. Distributions show clear separability of the two signals in this case. As the *E. coli* concentration increases, opsonization of the microbe remains at the maximum level and decreases only for high microbe centrations (greater than 10^14^ cells per litre, ratio of 20:1). Host opsonization nevertheless increases to a maximum of about 1000 molecules per cell. The signals still remain clearly separable until a concentration of 10^17^ *E. coli* cells per litre is reached. This corresponds to a ratio of 20000: 1 *E. coli* cells compared to erythrocytes. Even at this high ratio, separability can be achieved, but induces some autoreactivity, as can be seen from subfigure (V) in Fig. 8.

**Figure 8:**
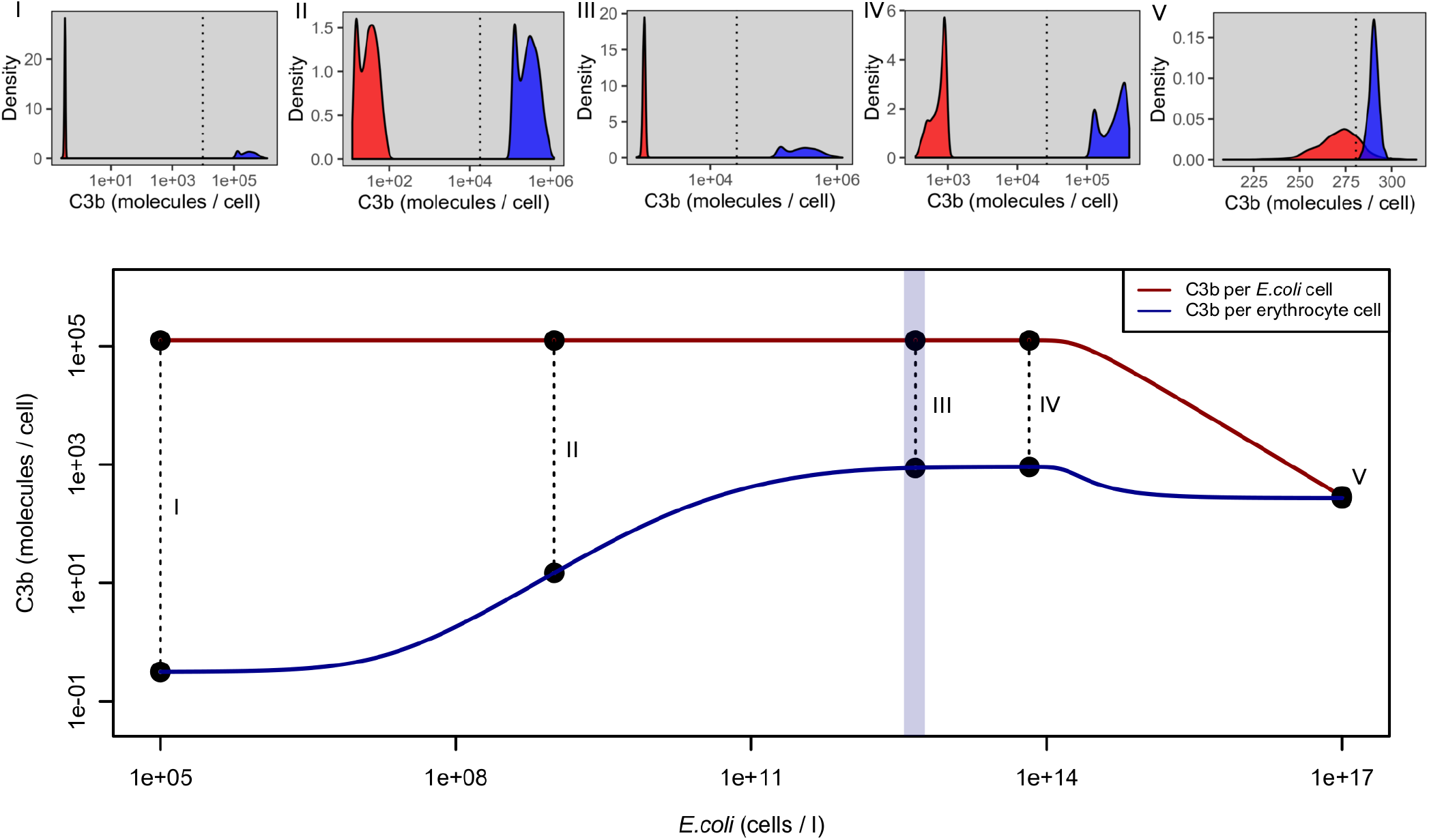
Computed mean C3b opsonization per cell if factor H cannot be acquired on the surface of the microbial cell. Erythrocyte counts used are indicated by the light blue shaded area (mean 4.64·10^12^ erythrocytes per litre, low: 1st percentile, high: 99th percentile, see Fig. A1). Microbial cell densities for subfigures (I)-(V) can be seen from the corresponding points in the lower panel. Curves represent mean C3b deposition on host and *E. coli* surfaces (double logarithmic plot). Subfigures represent C3b distributions at several *E. coli* concentrations. The dotted vertical lines in subfigures (I)-(V) represent the mean value of the optimal threshold interval to distinguish the two signals described in the section above. C3b opsonization remains low for host cells and high for microbial cells in a wide range of microbial cell densities and the signals are mostly well separable. Only for very large microbial cell densities they become inseparable because of insufficient production of C3 (see Fig. A3)

## Discussion

Here, we have analysed molecular crypsis by microbial pathogens involving human factor H. In this case, crypsis is defensive and aggressive at the same time because the mimics’ intention is invasion of unprotected host cells and mimicry makes them less suspicious, but at the same time other cell lines of the host can be regarded predators of the mimics so they need to hide from them in a defensive manner.

In particular, we established a mathematical model of molecular crypsis by *Candida, albicans*. It integrates methods from signal detection and protein interaction modelling. By choosing appropriate parameter values, it is easily adaptable to be applied to other pathogens. Results were compared to a strain of *Escherichia coli* that is considered not to perform molecular crypsis using FH.

Our results indicate that if microbes cannot acquire FH on their cell surface, the alternative pathway of complement is active on microbial cells and inactive on host cells, clearly separating self from non-self over a wide range of parameters. We demonstrated that, if pathogens are able to acquire FH on their surface, the effectiveness of complement may depend substantially and highly non-linearly on the quantity (and FH binding quality) of pathogen surface in the blood. This may come as a surprise because intuitively one would expect a linear dependence of opsonization on the target surface present in the medium.

The failure of molecular crypsis to be successful at low pathogen concentrations can be explained by the fact that, due to the positive feedback loop leading to amplification of C3B. there is no pure competition between C3B and FH. Rather, it is important which of the two binds first. This corresponds to the idea that at the very start of an infection, a single molecule of C3b encountering an unprotected surface may initiate a positive feedback loop of C3b generation (amplification) on that surface that is faster than the degradation or inactivation of C3b by surface-acquired FH. In contrast. C3b encountering a surface protected already by FH may not initiate C3b amplification. For increasing numbers of pathogen cells in the medium chances increase that a sufficiently high protection by FH is acquired on the cell surface before the first C3b molecule encounters this specific surface. This is because the production of activated C3b in the plasma is constant and shared by all cells in the medium.

This result is supported by experiments performed by Stone et al. (1974). They infused varying concentrations of 10^5^ to 10^19^ *C. albicans* cells per litre at a rate of 1 ml per minute into the portal vein of mongrel dogs and measured the presence of *C. albicans* in several compartments of the blood system and other tissues after 10 minutes (see Fig. 9). Their results correlate with our overall finding that clearance of mimetic pathogens is feasible at a low pathogen concentration, but not at a high concentration. For *C.albicans* concentrations below 10^11^ cells per litre, pathogen cells could only be detected at the infusion site and the liver. In subsequent parts of the blood system *C. albicans* was cleared successfully. For higher infusion rates nevertheless *C. albicans* was detected in all parts of the blood system. These results support our predicted transition point of the regime where molecular crypsis is not successful to the regime where it is successful, although the transition is not as sharp as predicted by our model. Note that we have not assumed, in each simulation, the *C. albicans* concentration to be time-dependent and that we have explicitly modelled the inflow of FH and C3 to regions different from the production site (the liver, see section A. 1.6).

**Figure 9:**
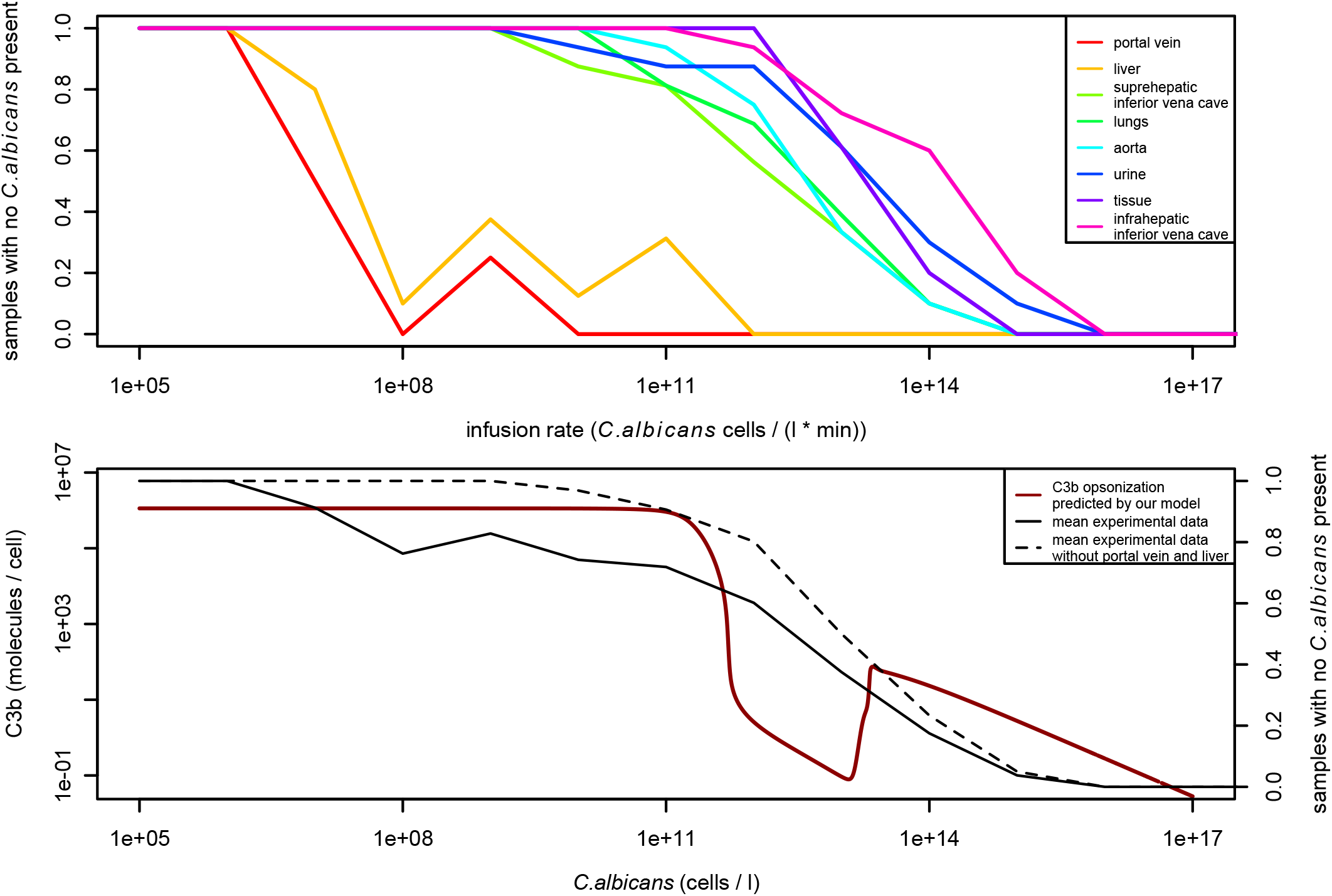
Effectiveness in clearing *C. albicans* cells from the blood of mongrel dogs based on varying infusion rates of *C. albicans* into the portal vein. Data was taken from Stone et al. (1974). Fig. 8. Data was converted from samples positive for *C. albicans* to relative effectiveness in clearing *C. albicans* cells and scaled to units used in our model for better comparability. Top: Samples with no *C. albicans* cells present after 10 minutes. measured in different compartments of the blood system. For infusion rates below 10^11^ cells per litre per minute, *C. albicans* cells are removed to a great deal by the liver and cannot be detected in subsequent compartments of the blood system. For higher infusion rates *C. albicans* cells are present in all compartments. Bottom: Comparison to results of our model. The predicted C3b opsonization (red line) correlates to the effectiveness in clearing *C. albicans* from the blood (black lines). High opsonization means clear identifìability of mimetic pathogens and therefore easy clearance. As the predicted opsonization decreases, also a decrease in clearance of *C. albicans* cells can be observed, especially in compartments subsequent to the liver (dashed line).

Autoreactivity at high infusion rates was mentioned by Stone et al. (1974), but not quantified, so the transition point to the regime where autoreactivity may occur could not be compared. Nevertheless in the experiments by Stone et al. (1974) no increase in *C. albicans* clearing effectiveness can be observed at around 10^14^ *C. albicans* cells per litre, indicating that the short increase in *C. albicans* opsonization to about 100 molecules per cell is not sufficient to activate complement.

From the above, we can conlude that the the positive feedback loop of C3B amplification is beneficial for host cells in several respects. It prevents molecular crypsis from being perfect at low pathogen load and decreases the response time. It allows the host to some extent to shape the environment.

Regarding spatiality. Pangburn and Müller-Eberhard (1984) showed that cells with activated complement and close proximity tend to “infect” each other with C3b. In our model this effect is only partially treated by increasing the rate of binding to the originating surface, but distributing the remaining increased C3b concentration to all cells in the medium (not only to cells in close proximity). If we neglect the scaling factor, we observe different results. indicating that spatiality really plays an important role (see Fig. A7). We suggest that simulations taking into account the spatial distribution and dynamics of *C. albicans* cells and erythrocytes (especially in differently-sized blood vessels, like veins and capillaries), would be very helpful to fully understand the effects of molecular crypsis by pathogenic microbes. For example, the concentration of the same amount of pathogens invading a capillary will be higher compared to veins, simply because the “local” volume of capillaries is smaller than for veins. Also blood viscosity varies with the radius of the vessel (Pries et al., 1992). Besides the different susceptibility of tissues to pathogen invasion, this could help to understand why several autoimmune diseases like age-related macular degeneration (AMD) seem to occur tissue-specific.

The effect of increasing opsonization on host cells but not on *C. albicans* cells at increasing pathogen concentrations can be explained by a higher binding affinity of FH to *C. albicans* surfaces than to host surfaces. The dissociation constant of FH at *C. albicans* surfaces is in the nanomolar range, while the dissociation constant of FH at host surfaces is in the micro-molar range. So the binding of FH to *C. albicans* surfaces is stronger than to host surfaces. This means that if we assume a somehow limited FH production. the pathogen can gain a substantial advantage in FH binding. As a result it is able to sequester dissociating protection from host cells (in addition to newly produced FH molecules), leaving them unprotected regarding C3b opsonization (see Fig. A4 and A9). Of course, the actual values will depend on the actual maximum expression rate of FH, which is hard to determine accurately (see Fig. A3 and A6).

The question remains whether the host can avoid perfect mimicry as an evasion mechanism of the pathogen. One option for the host would be to change the total amount of complement factors, which cannot be influenced by the pathogen. However, our model shows that changing these amounts does not help because the pathogen can always bind an appropriate fraction of factor H.

To respond to a decrease in the dissociation constant of FH by the pathogen, the host should do the same and could, for example, adapt the speed of C3b amplification while increasing binding affinities to FH, reaching the same balanced system but with an advantage in FH binding. Importantly, the alternative pathway of complement is not the only defence mechanism of the host. We can imagine a scenario following the so-called Red queen hypothesis (Brockhurst and Koskella. 2013). where (spatially related) hosts and pathogens evolve in a permanent arms race, trying to be always one step ahead of the other (Lively and Dybdahl, 2000). If one of them is not able to adapt to the advance of the other any more, it will lose the race. Maximizing binding affinities to FH nevertheless may become harder and harder in each round and is certainly limited by physical constraints and side-effects to other systems. This means that at some point, it could have been beneficial (or unavoidable) for the host to develop a completely new system of defence against mimicking pathogens.

Besides maximizing binding affinities, another way of gaining an advantage for hosts and pathogens, is the production of additional binding sites. For example C3đ. a cleavage product of C3b. can provide new binding sites for FH (Perkins et al., 2014). While it is generated on the surface of pathogens capable of molecular crypsis as well (since they also utilise FH to cleave C3b). it could be a mechanism of producing binding sites faster than the pathogen, if this cleavage product binds preferentially to host surfaces. Additionally this strategy practically does not involve any costs, since C3b is already needed to ensure complement activity and is cleaved on host cell surfaces anyway. The results of the population based model clearly indicated that both the host and the pathogen adjust the binding affinities and binding sites of factor H ligands to a maximum, although binding site production involves costs.

The fitness landscape of pathogens obtained by the evolutionary model is in line with results by Holen and Johnstone (2004), showing that costs of mimicry can lead to mimetic dimorphism (explaining why not all pathogens do mimicry). We observed two fitness maxima, one local maximum if no binding sites were produced at all and one global at perfect resemblance. This can be explained as follows: The benefit of mimicry is described by a constant basal growth rate minus a linear function of costs plus a saturation function above a threshold. The sum gives a non-linear function with two maxima. It does not pay at all to have a low degree of mimicry. For example, a fly being purely black or purely yellow has achieved half of the colouring of wasps but has (nearly) no mimicry effect. No mimicry is better than low mimicry, because the former is the local maximum at zero investment. The global maximum is achieved at a certain high investment (all or nothing). It is an interesting question how the global maximum could be attained during evolution, because it is difficult to explain by micro-evolution.

One way of avoiding perfect resemblance of pathogens in a population of hosts could be the utilization of polymorphisms of relevant proteins. On the host side there is the Y402H polymorphism in the FH gene and there is also a factor H like protein (FHL-1) which is an alternatively spliced product of the FH gene. It mainly consists of the first seven (of the 20) small consensus repeats (SCRs) responsible for binding. Those polymorphisms make adaptation to specific environments harder, as was already proposed by Damian (1964) in its initial study on molecular mimicry due to antigen sharing by hosts and parasites. There he suspected the ABO-polymorphism. defining the human blood groups, to be a consequence of molecular mimicry. On the pathogen side, there are also polymorphisms observable. For example pathogens often have more than one FH acquiring protein (GRASP) with slightly different binding affinity and different expression levels. Those could be adaptations to host polymorphisms, but also provide some robustness of the system if one receptor type gets lost or inefficient somehow.

It remains to be studied in more detail above which C3b concentration phagocytes decide to attack cells and if this threshold is adaptable. Based on methods of signalling theory, one can calculate optimal ways of discriminating two signals, as was shown in the Methods section. Still, it is hard to quantify the benefit of correctly attacking a pathogen and the cost of erroneously attacking a host cell. Generally we can assume none of the errors to be negligible, as was shown in a previous paper (Hummert et al., 2017). If we weight both errors equally, as was done in most of the simulations, we see that at high pathogen concentrations there is no optimal decision. The model would predict for this case that all cells present in the medium should be attacked. This is clearly not the optimal response of the host. In theory, this could only be prevented by zero benefits in attacking pathogen cells or equivalently infinite costs of attacking host cells. Such benefits and costs would be, on the other hand, non-optimal for low pathogen concentrations, as they may allow infections.

It would be optimal for the host to adapt its attack threshold based on the pathogen concentration in the blood. The threshold should be shifted towards effectiveness in clearing mimetic pathogens for low pathogen concentrations and shifted towards avoidance of autoreactivity for high pathogen concentrations (although the host has to rely on other mechanisms of the immune system then). Nevertheless it seems unlikely that such an adaptation of the threshold depending on the pathogen load can be realized. This is mainly because pathogens are capable of cryptic mimesis, meaning by definition that the host can hardly sense the concentration of pathogens. In view of the physiological opsonization state of the host (no pathogen is present), we see that the mean opsonization per cell is less than one molecule. So it seems likely that phagocytes attack even if a single molecule of C3b is sensed, although they do not scan the entire surface, so there should be more than one molecule on the surface for robust detection.

In addition of changing the threshold directly by alteration of the phagocyte’s predation strategy, also alteration of the the binding effectiveness of FH to surfaces maybe an option to control general reactiveness of complement. This is because FH removes deposited C3b from cell surfaces and thus higher binding affinity of FH to host surfaces results in better protection (calmed down complement) of host cells. On the other hand this onesided alteration of protection of host cells will likely result in adaptation of the FH binding effectiveness of the mimic by exactly the same amount, as we showed in the population based model. So basically both distributions would shift by the same amount in the same direction, which is then mathematically equivalent of changing the attack threshold, since it affects both errors equally. So although this may have no effect on seperability of host and pathogen cells it is a way of easily adapting general aggressiveness of complement without the need for alternations in the phagocyte’s predation strategy. So in fact the Y402H polymorphism could be an example of adaptation of the attack threshold, where the YY homozygote has a reduced complement reaction, while the HH homozygote may react more aggressively and the YH heterozygote is somewhere in the middle (Hummert et al., 2017).

Regardless of how the attack threshold is fixed exactly, we see that in the case where there is a high concentration of pathogens performing molecular crypsis, host opsonization can reach a point where autoreactivity seems nearly unavoidable. This may have severe consequences especially in situations where the adaptive immune system is suppressed. In such a scenario the pathogen load can hardly be regulated at all, possibly leading to severe malfunctions of the immune system, like sepsis. Lethal sepsis can indeed occur, for example, as a consequence of a disseminated candidiasis (Spellberg et al., 2005). Depending on the specific microbial environment, the host may accept the cost of a certain degree of autoreactivity in an early stage of infection to avoid pathogen concentration reaching the regime of successful crypsis. As mentioned above, there is a polymorphism Y402H in the human FH gene, where the H variant predisposes individuals to age-related macular degeneration (Hageman et al., 2005). It is an interesting question arising from the results of our model, if the H variant of the polymorphism on the other hand could act protective in the context of sepsis.

Returning to the analogy of the Captain of Koepenick (Zuckmayer, 1932), it is worth noting that to distinguish true from false policemen, also other signals are needed, such as correct commands. The deception by the Captain of Koepenick was so perfect because he had learned and used the army commands. To avoid a perfect camouflage, the human adaptive immune system is used in addition to the innate immune system to recognize antigens as further signals for discrimination. For molecular mimicry to be successful, pathogens need to deceive both systems.

## Acknowledgement

Financial support by the German Research Foundation (DFG) within the Jena School for Microbial Communication (grant no. DFG-GSC 214/2) and the Transregio 124 (FungiNet, projects B1, C4 and C6) is gratefully acknowledged.

## Author contributions

CG, CS, PFZ and SS designed the study. CG, SG and SNL established the models and conducted the data analysis. CS and PFZ performed the experiments. All wrote the manuscript. CS, PFZ and SS supervised the project.

## A Appendix

### A.l Supplementary Model Description

#### A.1.1 C3b diffusion in the blood

Taken from Zewđe et al. (2016).

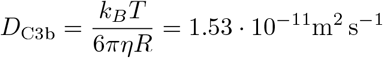

| Symbol | Value | Unit | Description |
| --- | --- | --- | --- |
| $k_B$ | $1.38 \cdot 10^{-23}$ | $\frac{\text{kgm}^2}{\text{s}^2\text{K}}$ | Boltzmann constant |
| $T$ | 310 | K | temperature |
| $\eta$ | 0.004 | $\frac{\text{kg}}{\text{m s}}$ | viscosity of blood |
| $R$ | $3.7 \cdot 10^{-9}$ | m | C3b radius |

#### A.1.2 Binding rate of reactive C3b

After proteolytic cleavage, C3b occurs as a short-lived reactive intermediate, called “nascent” C3b (nC3b) (Sim et al., 1981). nC3b is able to indiscriminately attach to different surfaces via an exposed internal thioester bond (Law et al., 1979; Tack et al., 1980; Law and Levine, 1977). For reactive C3b, we assume a diffusion controlled reaction (Collins and Kimball, 1949), where the binding reaction is assumed to occur spontaneously on contact with a cell surface.

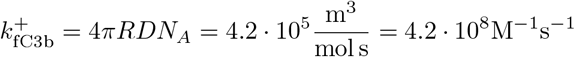

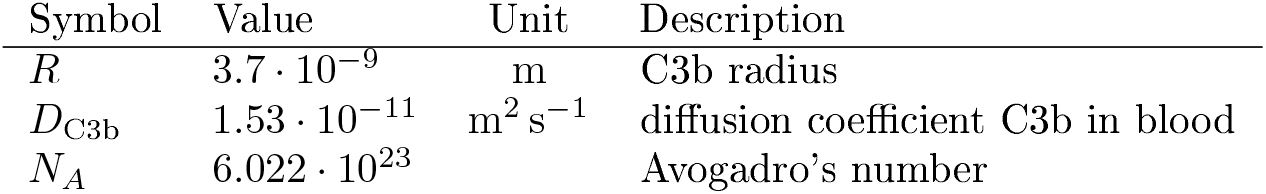

| Symbol | Value | Unit | Description |
| --- | --- | --- | --- |
| $R$ | $3.7 \cdot 10^{-9}$ | m | C3b radius |
| $D_{C3b}$ | $1.53 \cdot 10^{-11}$ | $\text{m}^2 \text{s}^{-1}$ | diffusion coefficient C3b in blood |
| $N_A$ | $6.022 \cdot 10^{23}$ | | Avogadro's number |

#### A.1.3 Decay rate of reactive C3b

Sim et al. (1981) measured a half live *t*_1/2_ of about 60 μs for nC3b to remain active, before it reacts with water to form fluiđ-phase C3b (fC3b). In the model, this process is described with the reaction nC3b → fC3b and the following rate:

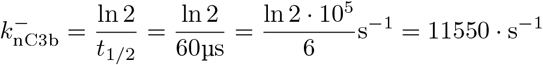

#### A.1.4 C3b active hemispheric region

With the half-time of C3B of

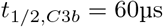

we obtain the time after which 90 % of C3b have been inactivated:

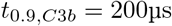

So reactive C3b may diffuse a distance of

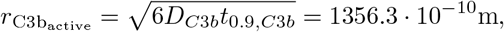

before 90 % of its activity is lost.

#### A.1.5 Scaling factor for the affinity of surface derived nascent C3b to the originating surface

Adapted from Zewde et al. (2016).

Although there is no discrimination in binding affinity of nC3b to pathogen or host surfaces, the spatial location of the C3b amplification unit C3bBb (production side) has to be considered. The binding affinity 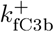 of free C3b will only hold in a well mixed environment, namely for nfC3b produced in fluid by a fC3bBb amplification unit. For surface bound C3bBb units (host hC3bBb or pathogen pC3bBb) the local concentration of surface will be different than assumed in a homogeneous environment (where surface does not actually exist, but is assumed a homogeneous concentration of binding sites in the medium).

Since we know the radius of 90 % activity of C3b (subsection A. 1.4), we can neglect the non-planarity of the cell surface in this region and calculate the binding sites surrounding a single hC3bBb or pC3bBb unit as

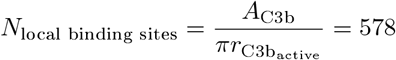

where *A*_c3b_ = 100 × 100Å = 10^-16^m^2^ is the surface area occupied by a single C3b molecule.

The local concentration of binding sites is then (assuming half spherical volume, because the membrane is impermeable to C3b)

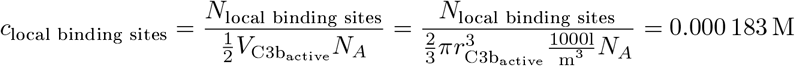

where *N_A_* = 6.022 · 10^23^ is Avogadro’s number.

To account for this in the model, we could multiply the global binding site concentration *c*_global binding sites_ of host and pathogen with the following factor:

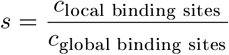

But since the reaction is assumed to follow irreversible mass-action kinetics, we can equivalently multiply host and pathogen nC3b binding affinities with the same factor:

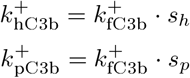

In summary this makes the binding affinity of nascent C3b to the originating surface higher than to other surfaces or fluid nascent C3b.

#### A.1.6 Inflow and outflow of relevant complement factors C3 and H in the blood stream

Complement factors were first treated as external metabolites in the model. But for very high binding site concentrations this led to unrealistic scenarios where the amplification of C3b on pathogen surfaces (consuming C3) affected fluid phase amplification (also consuming C3) in a way that an explosion of concentrations occurred. Therefore we decided to model the inflow and outflow rates of the relevant complement factors H and C3 explicitly. Actual expression rates are hard to determine and may be extensively dependant on consumption rates due to gene regulation networks. Therefore we simplified this process by the idea that we have a static production site (e.g. the liver), where the concentration of complement factors is always kept constant (physiological conditions obtained from the literature, see Table A1). It is only relevant then how often we pass this region based on mean bloođ-flow (cardiac output *Q*, blood volume *V* and the respective mean concent ration of H_*global*_ and C3_*global*_ at the production site):

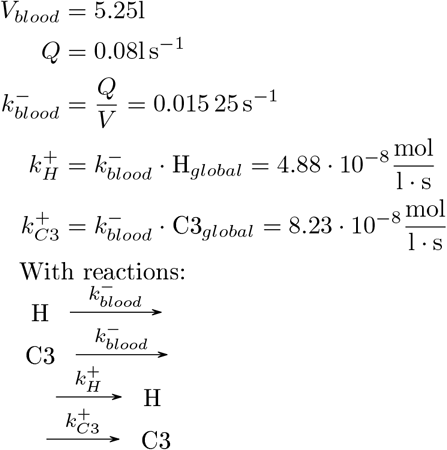

**Table A1:** Complement protein concentrations used in the model, as proposed by Zewde et al. (2016).

| Complement proteins | Concentration ($\mu\text{M}$) | Source |
| --- | --- | --- |
| C3 | 5.4 | Zipfel and Skerka (2015) |
| Factor H | 3.2 | Zipfel and Skerka (2015) |
| Factor B | 2.2 | Zipfel and Skerka (2015) |
| Factor D | 0.083 | Zipfel and Skerka (2015) |
| Factor I | 0.4 | Zipfel and Skerka (2015) |

#### A.1.7 Surface area distributions and erythrocyte count distribution used for sampling

**Figure A1:**
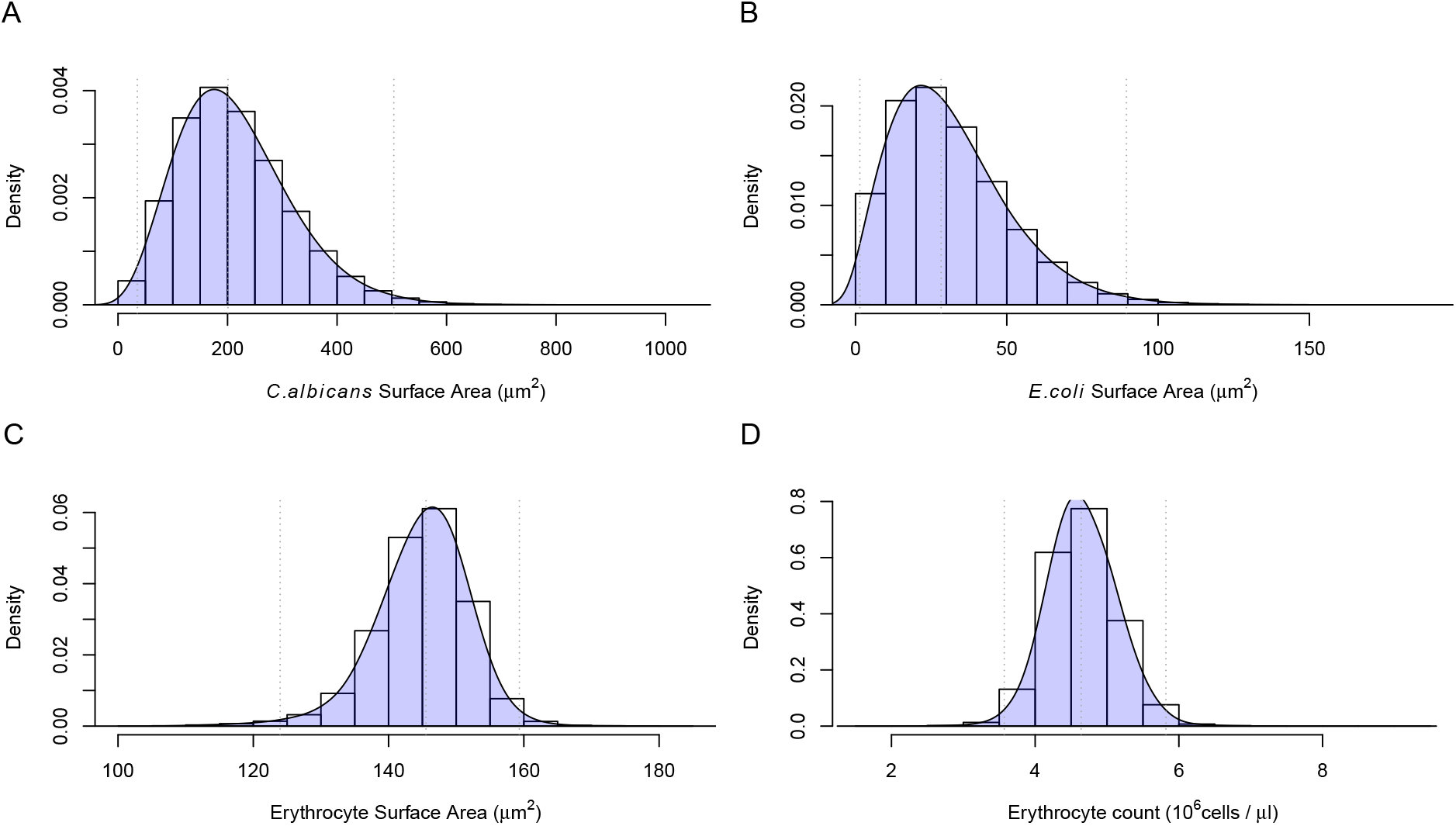
Surface area distributions and erythrocyte count distribution used for sampling. Dotted vertical lines represent the 1st and 99th percentiles of the respective densities. (A) Density of *C. albicans* surface area assuming a spherical shape with a normally distributed diameter (*μ* = 8μm, *σ* = 2μm). (B) Density of *E. coli* surface area assuming a spherical shape with a normally distributed diameter (*μ* = 3μm, *σ* = 1μm). Erythrocyte surface area (C) and count (D) were estimated from the combined NHANES datasets from 2001 to 2014 (Centers for Disease Control and Prevention (CDC). National Center for Health Statistics (NCHS). 2014). The erythrocyte count is given directly in the data. To estimate erythrocyte surface area we used the mean cell volume and assumed a cylindrical geometry with a radius-to-height ratio of 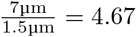 (based on mean values in the literature).

#### A.1.8 Reaction Diagram

**Figure A2:**
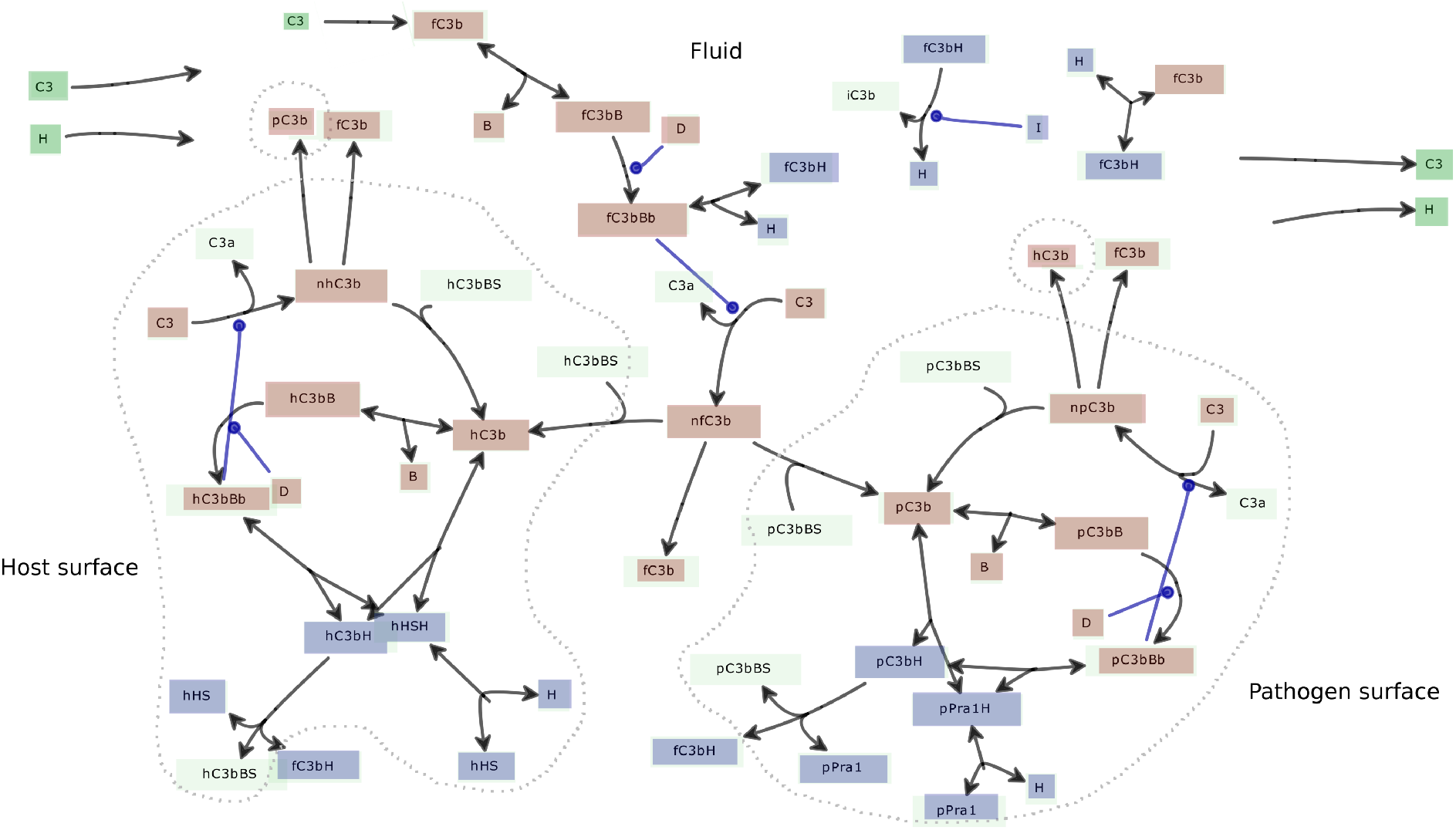
Diagram of all reactions occurring in the *C. albicans* model. Inflow and outflow are indicated in green, amplification in red and regulation in blue. Starting with the tick-over reaction of C3 into fC3b (top-left), fC3b next associates in fluid with factor B to form the fluid C3 proconvertase (fC3bB). fC3bB is activated by factor D to form the active fluid C3 convertase (fC3bBb). fC3bBb may associate with factor H to form fC3bH, which is cleaved into iC3b by the fluid degradation reactions using factor I (top-right reactions, indicated in blue). If not inactivated, the active fluid C3 convertase (fC3bBb) may convert C3 into C3a and fluid nascent C3b (fnC3b). fnC3b may attach to either host or pathogen surfaces (see section A. 1.2), or associate with water to form fC3b (see section A. 1.3), which might start the fluid amplification loop again (if not regulated). If fnC3b hits a surface, the same reactions as in fluid may occur, eventually leading to the formation of the active C3 convertase bound to surfaces (pC3bBb and hC3bBb). See also Figs. 1 and 3 in the main text for a schematic visualization of the reactions. The only differences in the reactions on surfaces and in fluid are that factor H may not attach to surfaces unless it is actively acquired by surface proteins (which are pPral for *C. albicans* and hHS for erythrocytes) and that a host or pathogen C3b binding site (hC3bBS and pC3bBS) is required for fnC3b to attach to surfaces.

#### A.1.9 Complement protein concentrations and Kinetic rate constants used in the model

**Table A2:** Kinetic rate constants used in the model, as proposed by Zewde et al. (2016).

| Biochemical reaction | Rate Constant | Value | Source |
| --- | --- | --- | --- |
| Hydrolysis of $\text{C3}(\text{H}_2\text{O})$ | $k_{\text{C3H}_2\text{O}}^+$ | $8.3 \times 10^{-10} \text{ ms}^{-1}$ | Pangburn et al. (1981) |
| Association of factor B to C3b | $k_{\text{C3bB}}^+$ | $0.000\,213 \mu\text{M}^{-1} \text{ ms}^{-1}$ | Chen et al. (2010) |
| Dissociation of complex C3bB | $k_{\text{C3bB}}^-$ | $0.000\,155 \text{ ms}^{-1}$ | Chen et al. (2010) |
| Attachment of fnC3b to host and pathogen | $k_{\text{fC3b}}^+$ | $0.42 \mu\text{M}^{-1} \text{ ms}^{-1}$ | Calculated, see A.1.2 |
| Attachment of hnC3b to host | $k_{\text{hC3b}}^+$ | varying | Calculated, see A.1.5 |
| Attachment of pnC3b to pathogen | $k_{\text{pC3b}}^+$ | varying | Calculated, see A.1.5 |
| Association of nC3b (fnC3b, hnC3b and pnC3b) to water | $k_{\text{nC3b}}^-$ | $11.55 \text{ ms}^{-1}$ | Calculated, see A.1.3 |
| Association of factor H to C3b | $k_{\text{C3bH}}^+$ | $0.0052 \mu\text{M}^{-1} \text{ ms}^{-1}$ | Pangburn and Mueller-Eberhard (1983) |
| Dissociation of complex C3bH | $k_{\text{C3bH}}^-$ | $0.0325 \text{ ms}^{-1}$ | Pangburn and Mueller-Eberhard (1983) |
| Association of factor H to heparin dp32/dp36 (HS) | $k_{\text{HSH}}^+$ | $0.0065 \mu\text{M}^{-1} \text{ ms}^{-1}$ | Perkins et al. (2014), estimated from dissociation constant |
| Dissociation of complex HSH | $k_{\text{HSH}}^-$ | $0.003\,25 \text{ ms}^{-1}$ | Perkins et al. (2014), estimated from dissociation constant |
| Association of factor H to Pra1 | $k_{\text{Pra1H}}^+$ | $0.8673 \mu\text{M}^{-1} \text{ ms}^{-1}$ | own measurements, estimated from dissociation constant |
| Dissociation of complex Pra1H | $k_{\text{Pra1H}}^-$ | $0.001\,62 \text{ ms}^{-1}$ | own measurements, estimated from dissociation constant |
| Inflow of factor H | $k_{\text{H}}^+$ | $4.88 \times 10^{-5} \mu\text{M} \text{ ms}^{-1}$ | Calculated, see A.1.6 |
| Inflow of C3 | $k_{\text{C3}}^+$ | $8.23 \times 10^{-5} \mu\text{M} \text{ ms}^{-1}$ | Calculated, see A.1.6 |
| Outflow of FH and C3 | $k_{\text{blood}}^-$ | $1.525 \times 10^{-5} \text{ ms}^{-1}$ | Calculated, see A.1.6 |
| Activation of complex C3bB by Factor D | $k_{\text{cat}}^{\text{C3bB}}$<br>$K_M^{\text{C3bB}}$ | $0.0021 \text{ ms}^{-1}$<br>$0.1 \mu\text{M}$ | Zewde et al. (2016) |
| Cleavage of C3 by C3 convertase, C3bBb | $k_{\text{cat}}^{\text{C3bBb}}$<br>$K_M^{\text{C3bBb}}$ | $0.0018 \text{ ms}^{-1}$<br>$5.9 \mu\text{M}$ | Pangburn and Müller-Eberhard (1986) |
| Cleavage of C3b by inhibitor Factor I | $k_{\text{cat}}^{\text{C3bH}}$<br>$K_M^{\text{C3bH}}$ | $0.0013 \text{ ms}^{-1}$<br>$0.25 \mu\text{M}$ | Pangburn and Mueller-Eberhard (1983) |

### A.2 Supplementary results and example dynamics

**Figure A3:**
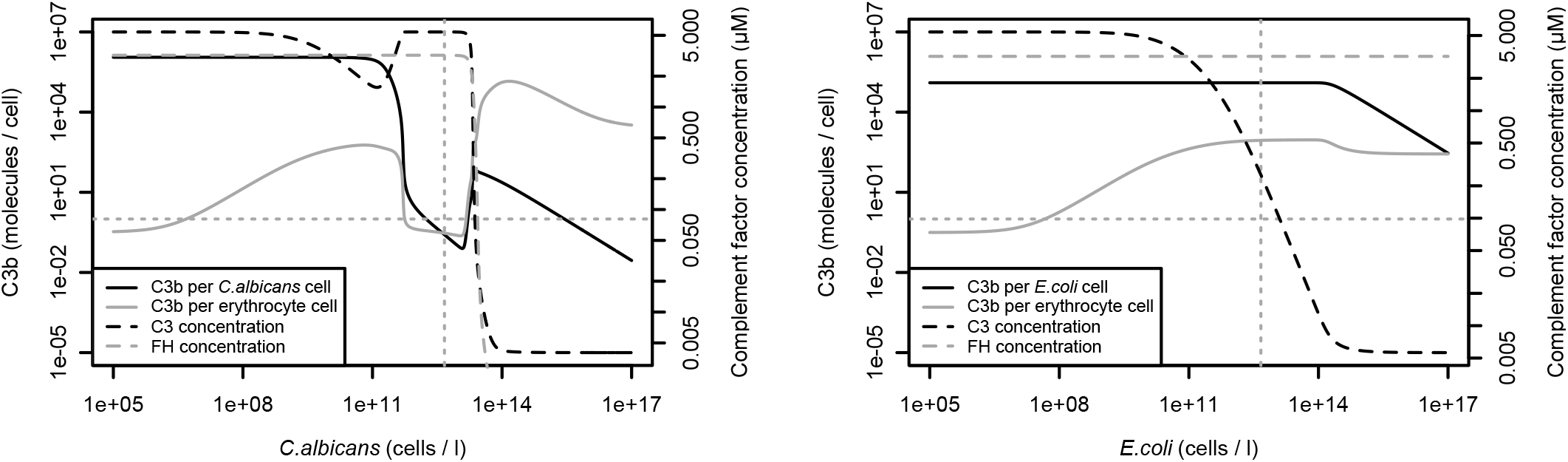
Opsonization states and relevant complement factor concentrations assuming inflow of C3 and FH limited only by blood flow. Same data as used for creating the figures of the main text. (Figs. 6 and img.opsonizatioriEColi). Left. *C. albicans*: after the phase of successful crypsis. complement proteins FH and C3 decrease drastically. FH drops faster and the concentration is insufficient to protect both species. Since the binding affinity to FH is stronger on *C. albicans* surfaces, host opsonization increases faster. Right. *E. coli:* FH concentrations are not affected by increasing pathogen concentrations, but for high *E. coli* concentrations. C3 production is insufficient to ensure opsonization of both species.

**Figure A4:**
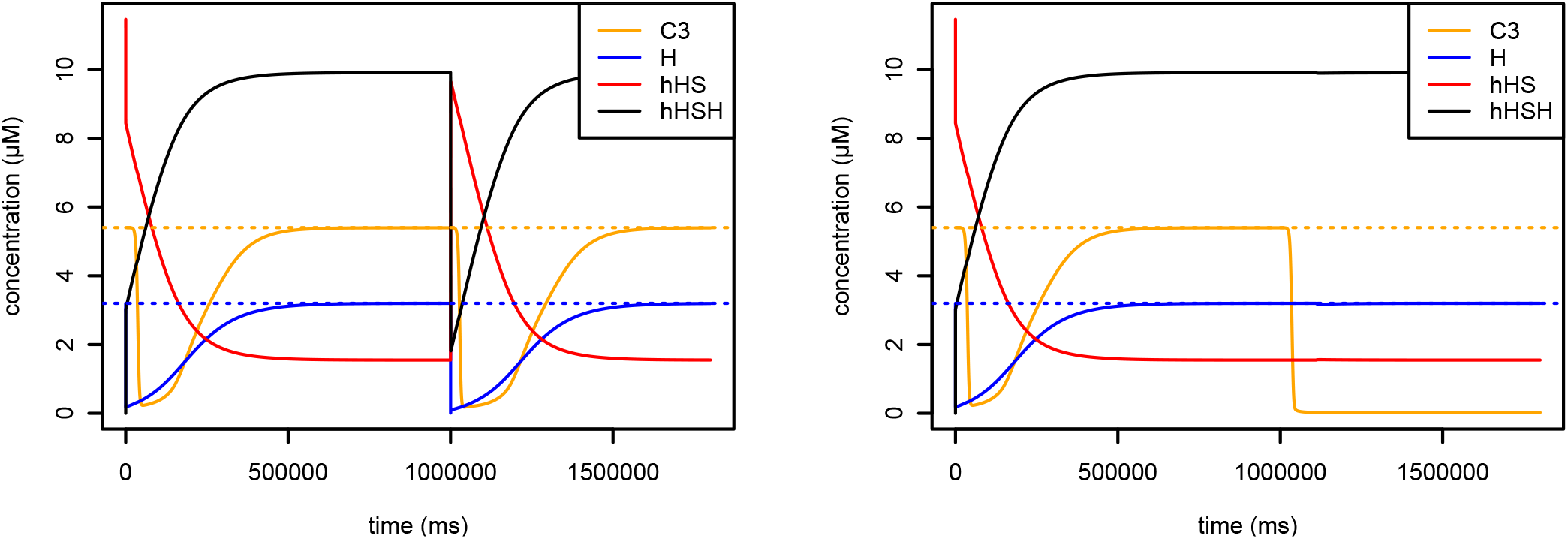
Example dynamics assuming inflow of C3 and FH limited only by blood flow. Cell concentrations were chosen equally for host and pathogen. Left. *C. albicans:* directly after pathogen arrival at 1000 seconds, host cells loose a great deal of FH protection within a few milliseconds, in which the free heparan sulfate concentration (hHS) increases, and the bound heparan sulfate concentration (hHSH) decreases. After a period of about 250-500 sec. the original state is restored. This is because the binding of FH to pathogen surfaces is much stronger and dissociating FH from host surfaces gets sequestered by *C. albicans* immediately (*C. albicans* Pral and PralH concentrations behave inversely, data not shown). Host protection recovers due to the inflow of FH. At even higher pathogen concentrations. FH inflow would not be sufficient to recover the protected state. In the case of *E. coli* (right) host protection is nearly unaffected, but C3 concentration decreases rapidly due to opsonization of the *E. coli* cells.

### A.3 Supplementary results and example dynamics using alternative assumptions

**Figure A5:**
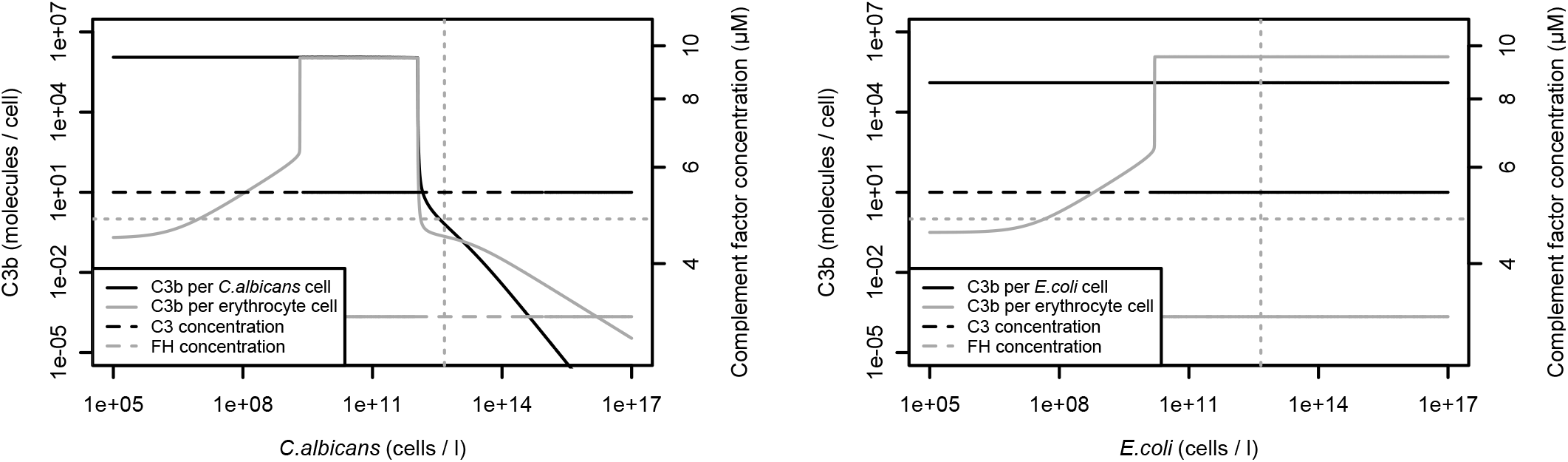
Opsonization states and relevant complement factor concentrations assuming unlimited inflow of C3 and FH. We see rapid opsonization of both species even for low pathogen concentrations. This is due to the fact that the production of nascent C3b on surfaces influences fluid phase C3b amplification in a way that the regulation by FH is not sufficient to prevent amplification. Since C3 inflow is not limited, fluid phase nascent C3b explodes to infinite concentrations. A short decrease in C3 concentration (due to limited production / inflow) after complement activation seems necessary to prevent this. It is interesting to note nevertheless, that in the case of unlimited C3 and FH production, opsonization at high *C. albicans* (left) concentrations decreases for host and pathogen. This means that autoreactivity is unlikely to occur and local expression of FH. as it is done by for example epithelial cells, may help to prevent autoreactivity at least to some degree (expression must have certain limits).

**Figure A6:**
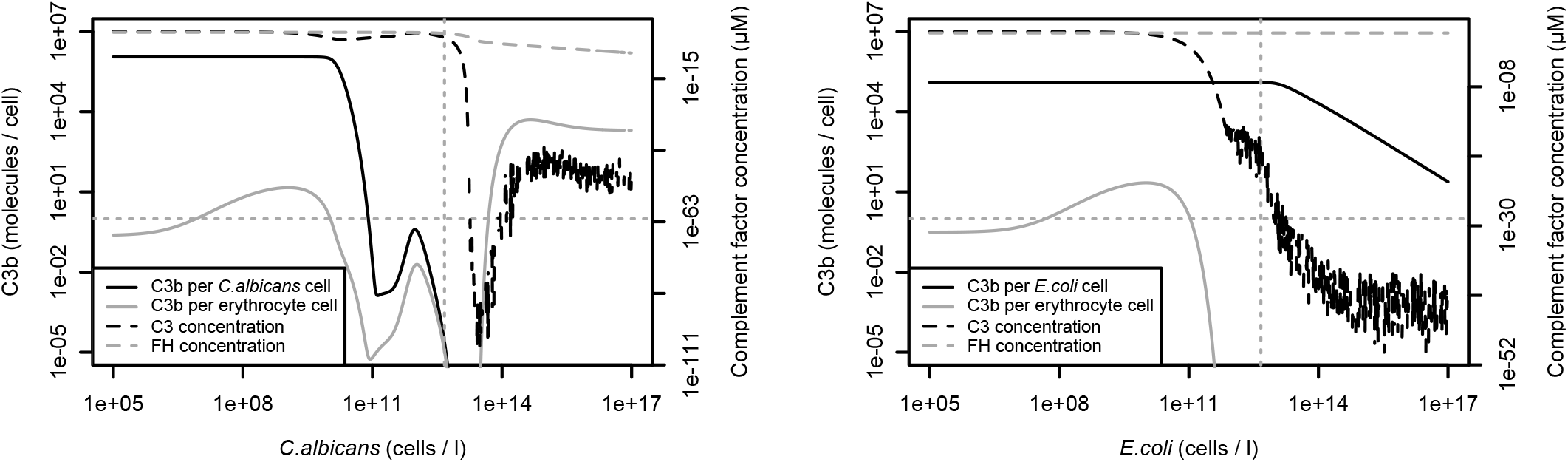
Opsonization states and relevant complement factor concentrations assuming no inflow of C3 and FH. We see a faster drop in opsonization if molecular crypsis can be performed (left) but a short increase afterwards, before opsonization reaches zero. For high *C. albicans* concentrations, the pathogen remains without opsonization at all and only the host is opsonized. *E. coli* (right) opsonization is similar to the case with inflow, but the host has zero C3b bound on surfaces. Note that complement concentrations are very small for high pathogen concentrations and the numerical precision is not sufficient to perform accurate simulations.

**Figure A7:**
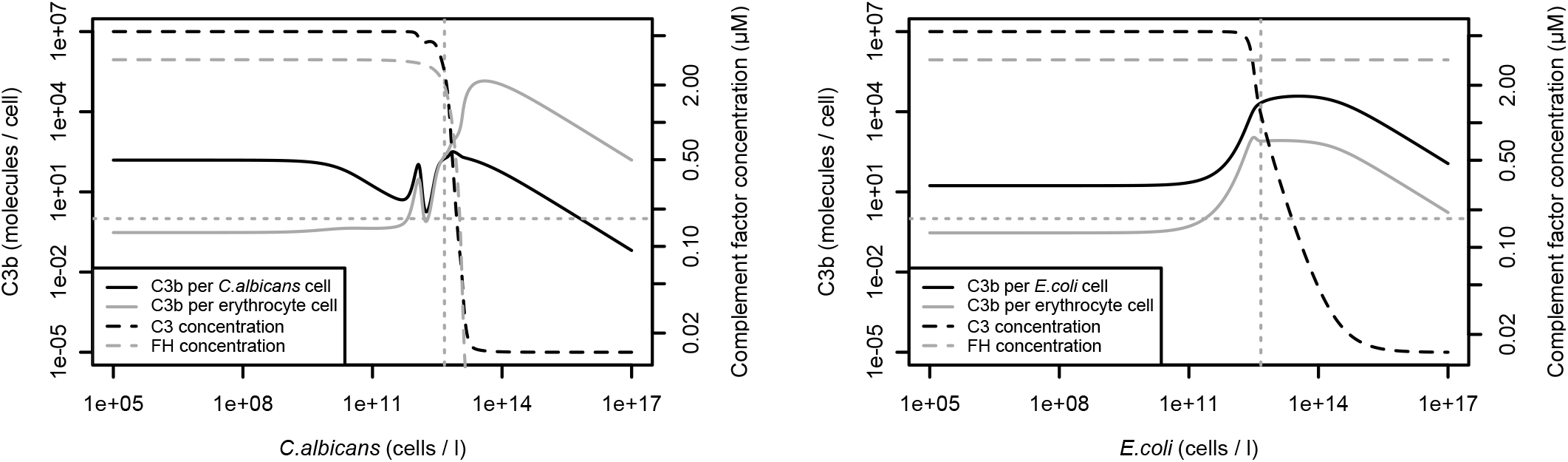
Opsonization states and relevant complement factor concentrations without the scaling factor accounting for spatial effects. Without increased binding affinity of surface derived C3b to its originating surface, we see the same qualitative behaviour, but with less clear separability between host and pathogen cells. The regime where molecular crypsis is successful (left) is shorter and a oscillation of opsonization occurs. For *E. coli* (right) there could be a phase of inseparability even if the pathogen cells are less abundant than the host cells.

**Figure A8:**
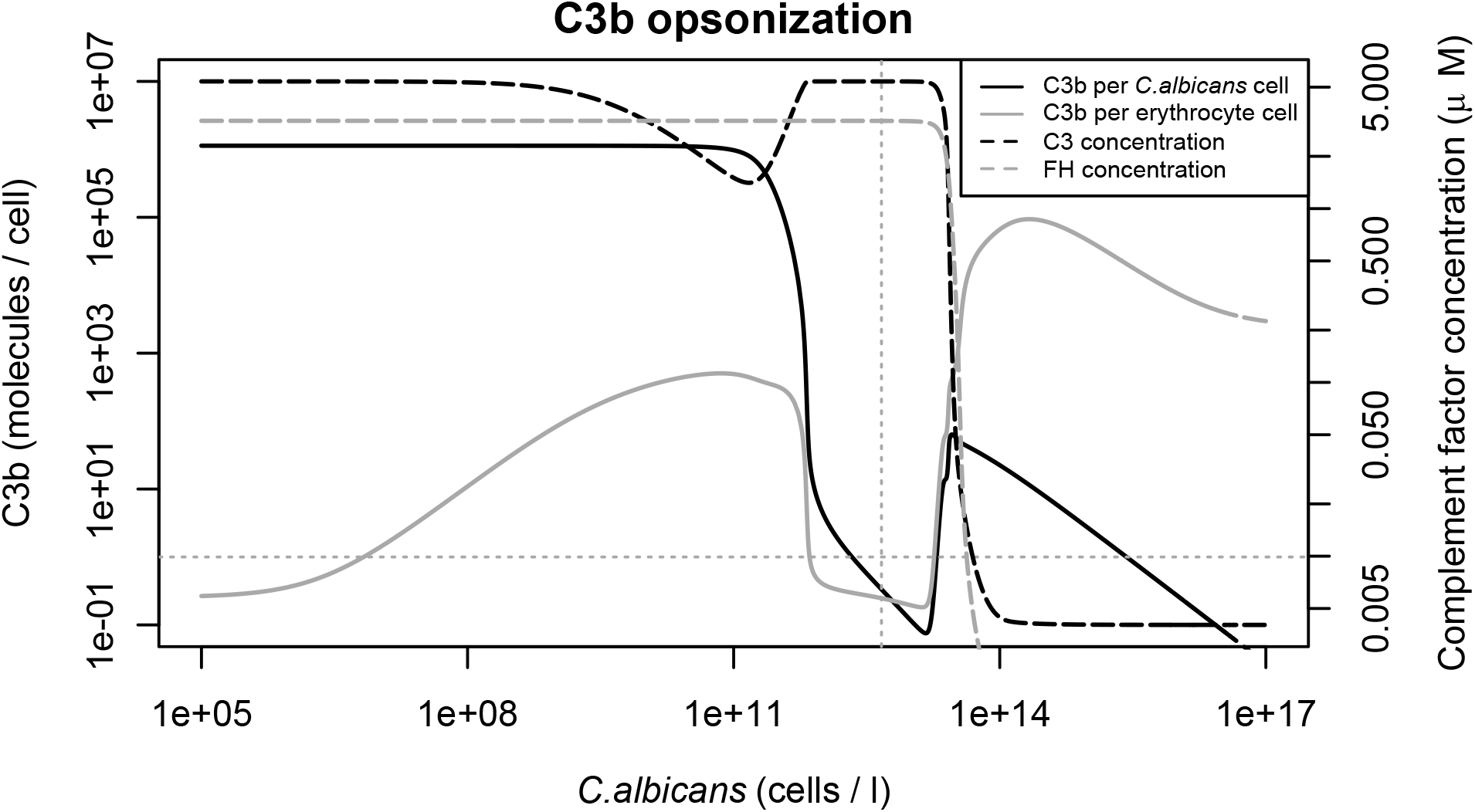
Opsonization states and relevant complement factor concentrations with higher heparan sulfate concentration and lower Pral concentration on the surfaces. Heparan sulfate and Pra1 were increased and decreased, respectively, by 50 *%* compared to the standard values used. Variation in FH binding sites does not alter the dynamics in general, but a higher concentration of *C. albicans* cells compared to erythrocytes is needed to achieve the same qualitative behaviour as in Figs. 6 and A3.

**Figure A9:**
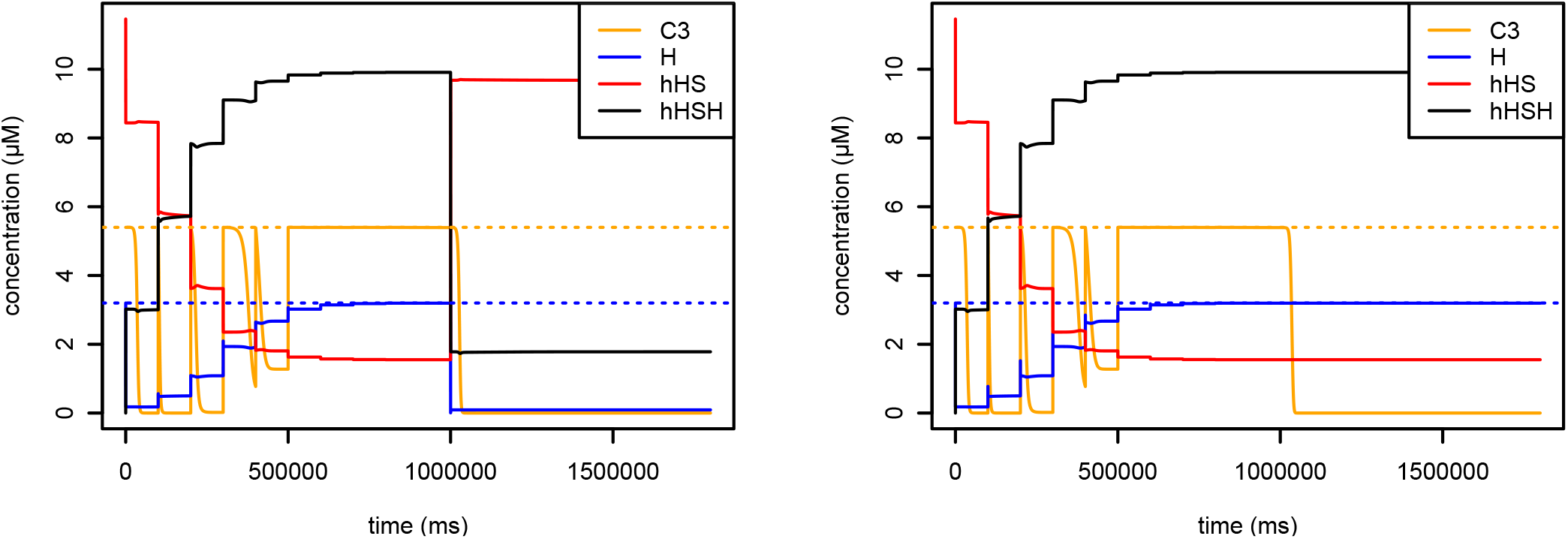
Example dynamics assuming no inflow of C3 and FH. Cell concentrations were chosen equally for host and pathogen. Basically the same behaviour as in Fig. A4. but the host cannot recover its protected state and may be opsonized even for equal host and pathogen cell densities (at least if enough C3 is present).

### A.4 Complete ODE system

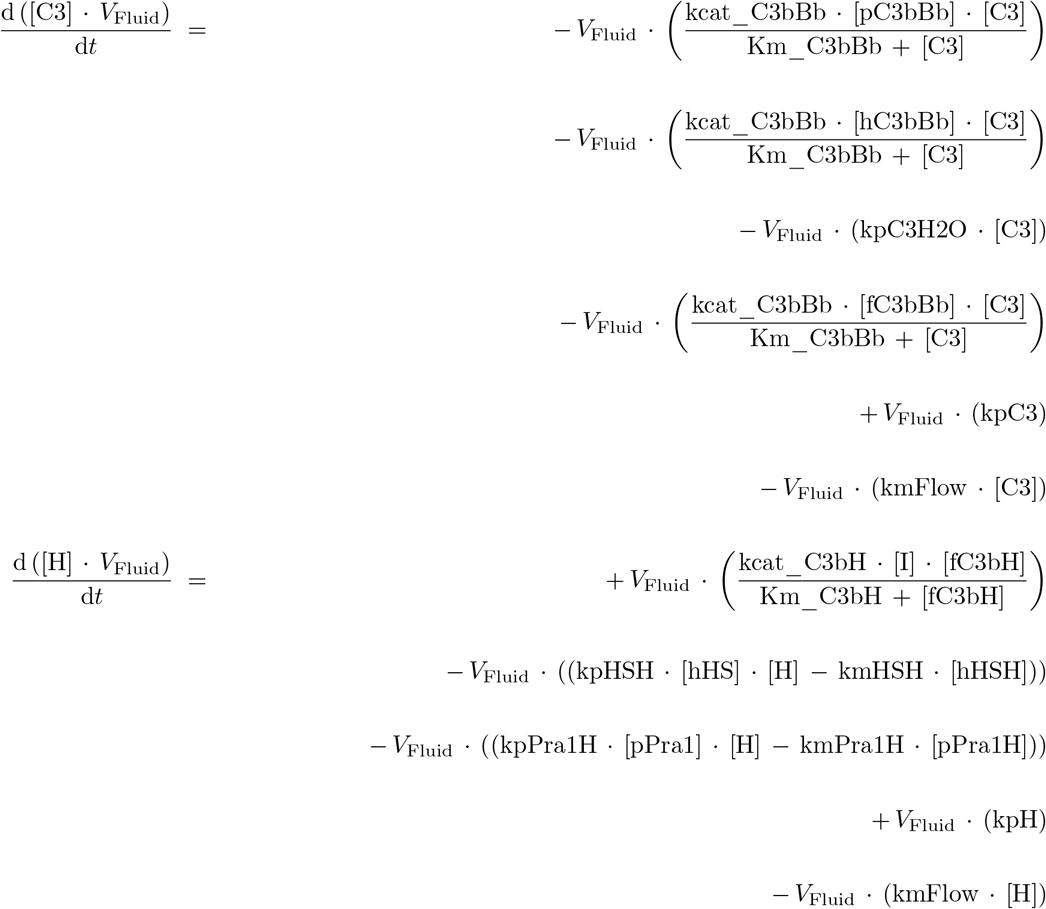

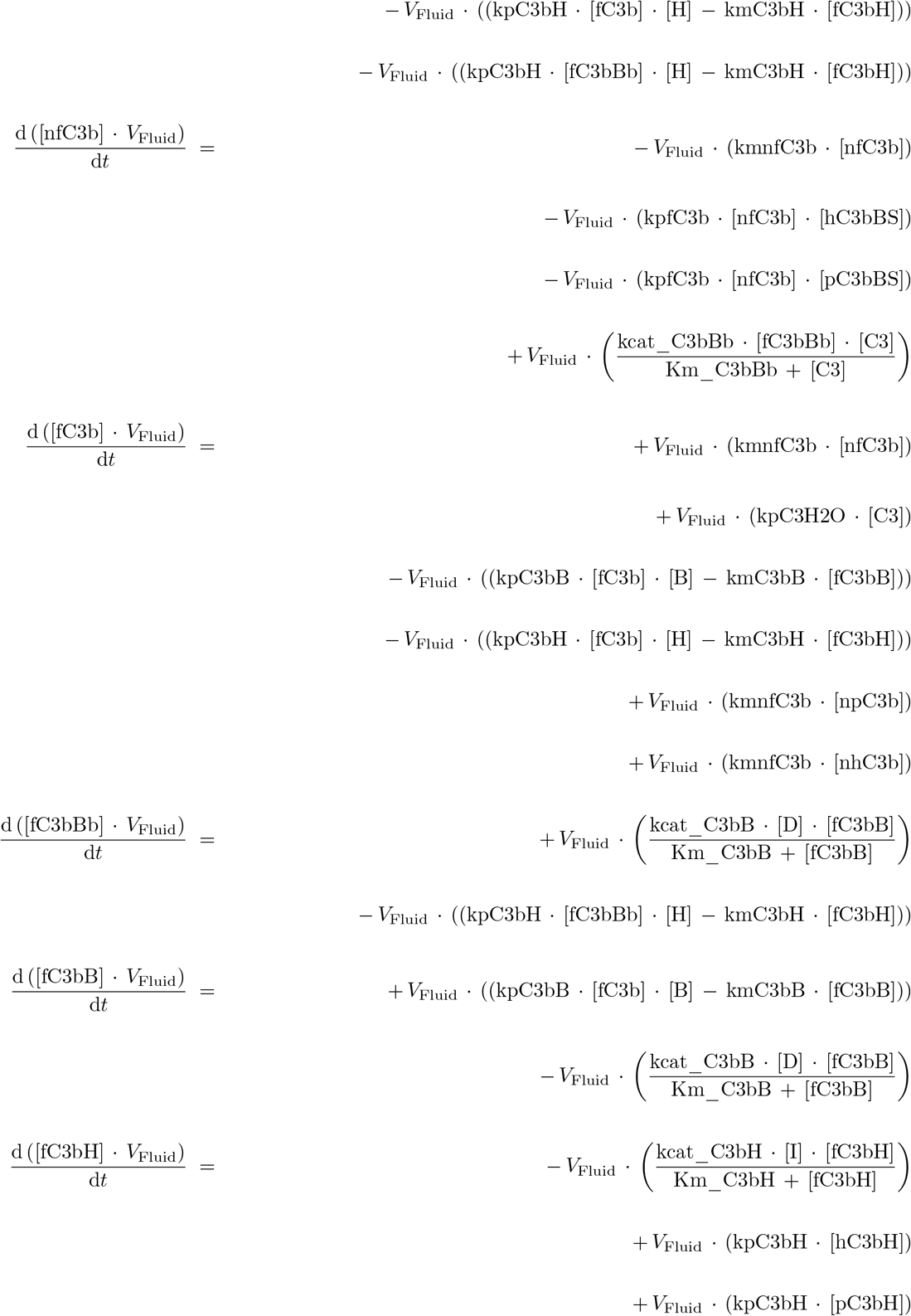

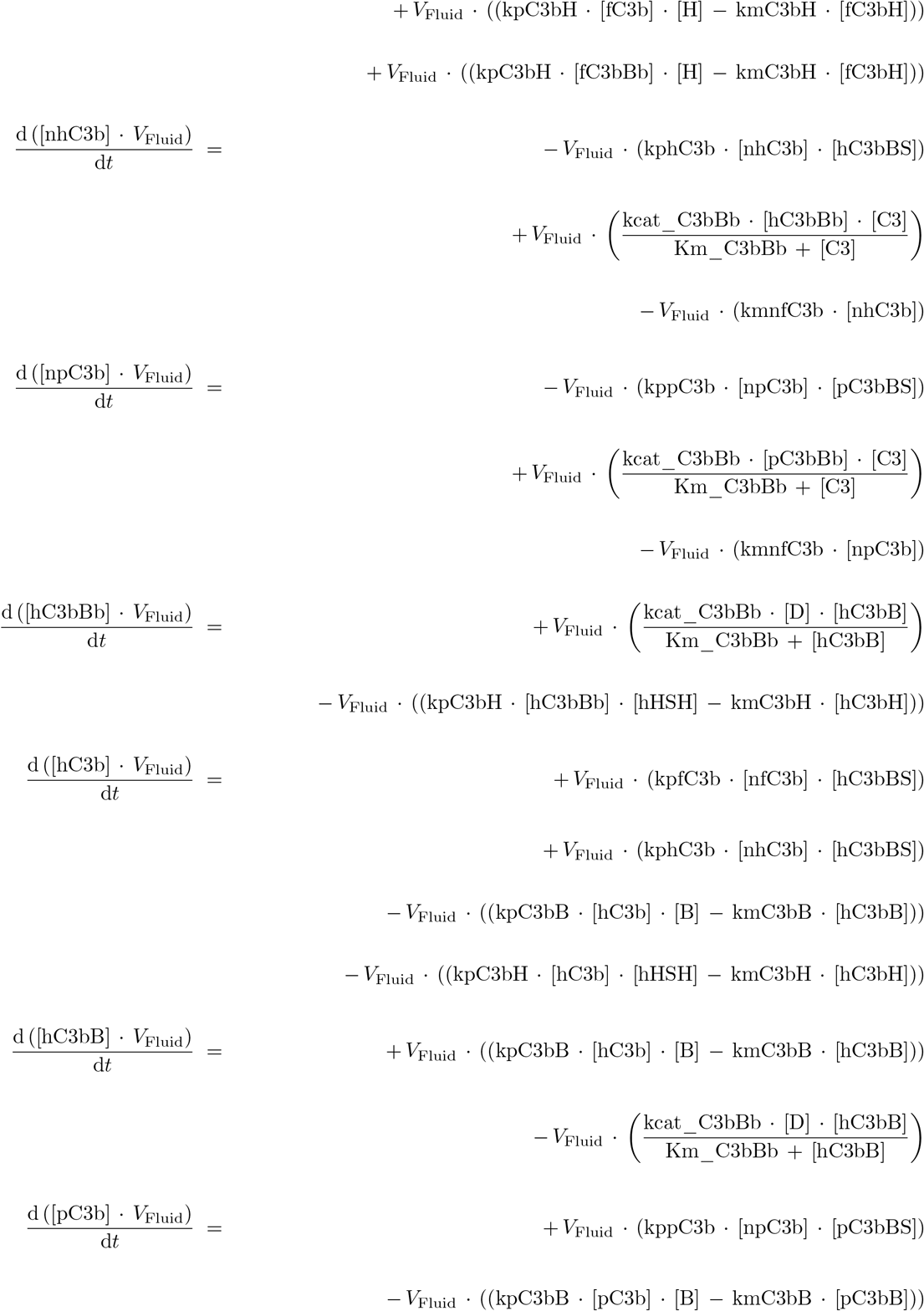

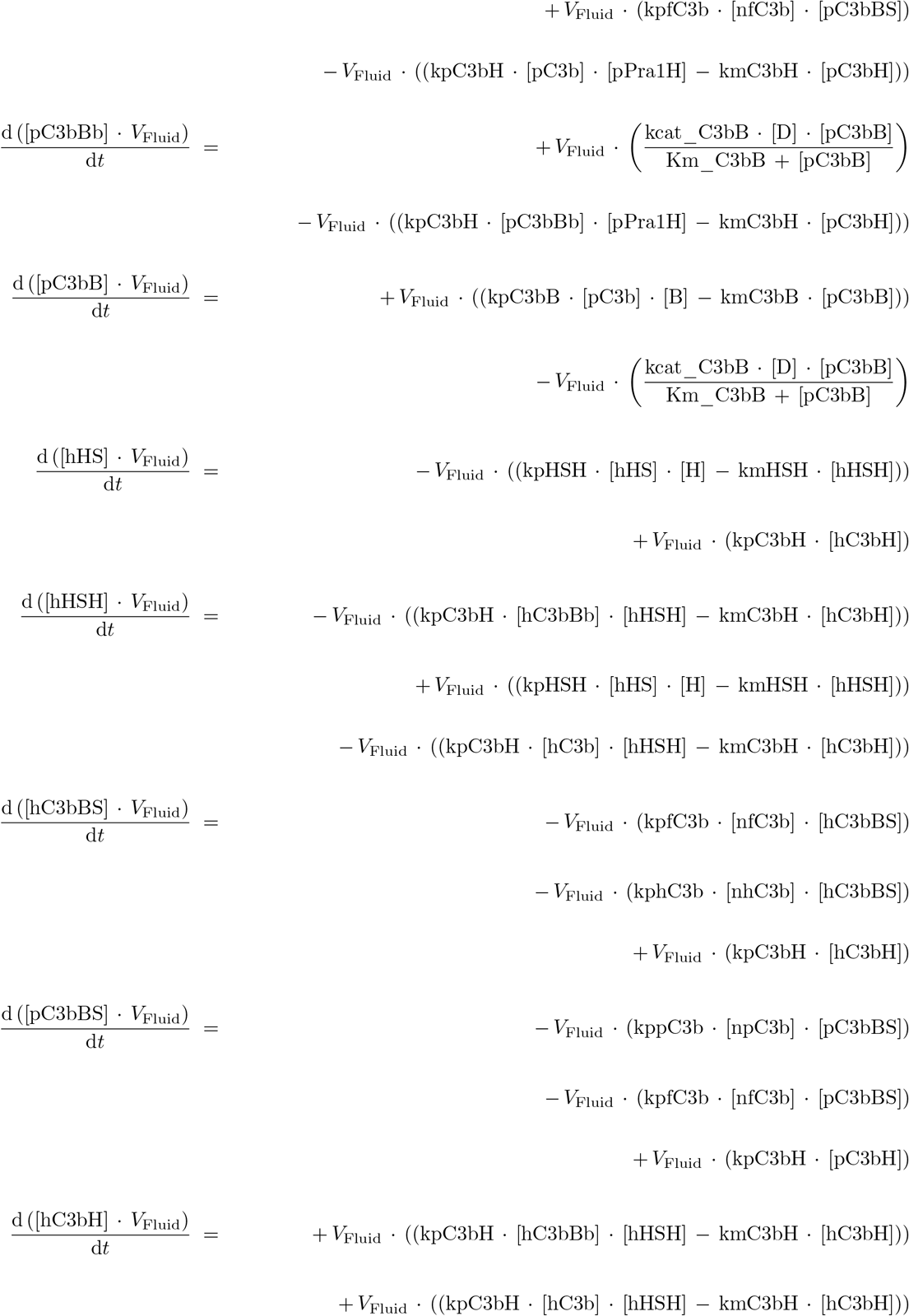

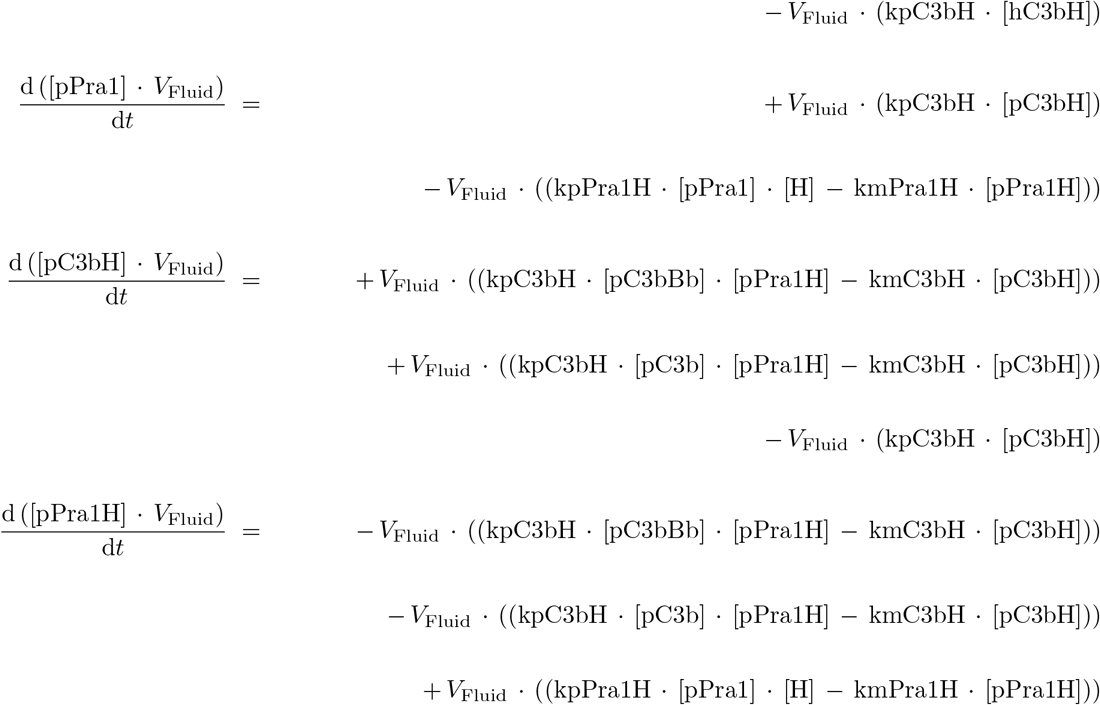

